# Recurrent hybridization and gene flow shaped Norway and Siberian spruce evolutionary history over multiple glacial cycles

**DOI:** 10.1101/2023.10.04.560811

**Authors:** Qiujie Zhou, Piyal Karunarathne, Lili Andersson-Li, Chen Chen, Lars Opgenoorth, Katrin Heer, Giovanni Giuseppe Vendramin, Andrea Piotti, Elena Nakvasina, Martin Lascoux, Pascal Milesi

## Abstract

Over the last decades, extensive genome-wide resequencing studies have highlighted the extent of hybridization and introgression between closely related species. Animal and plant species went through cycles of contractions and expansions as a result of glacial cycles. These repeated sequences of reproductive isolation and admixture at continental scales have led to the accumulation over time of an ancient, deep-seated and complex genetic structure. This structure was blurred by extensive gene flow, or reinforced by strong local adaptation. This already multi-layered structure has often been further enhanced by hybridization.

We investigated this complexity in Norway spruce (*Picea abies*) and Siberian spruce (*P. obovata*), two closely related species dominating Eurasian boreal forests and forming a vast hybrid zone. Here, we genotyped 542 individuals of both species and their hybrids at 480K SNPs. Individuals came from 55 populations, extending from western Europe to Siberia with a focus on the main hybrid zone. Despite extensive gene flow and a clear Isolation-by-Distance pattern at the continental scale, distinct genetic clusters emerged, indicating barriers and corridors to migration. Coalescent-based demographic inferences revealed that Norway and Siberian spruce repeatedly hybridized during the Pleistocene with introgression pattern varying depending on the latitude. In northern ranges, *P. obovata* expanded into *P. abies* while *P. abies* expanded into *P. obovata* in the southern parts. Two cryptic refugia located in the large hybrid zone played a critical role in shaping the current distribution of the two species. Our study highlights the importance of considering the whole species complex instead of separate entities to shed light on their complex demographic histories.

## Introduction

Both animal and plant species went through cycles of contractions and expansions as a result of glacial cycles (Fletcher et al. 2010; Nevado et al. 2018; Shuvaev et al. 2023). These cycles were generally accompanied by sequences of reproductive isolation and strong genetic divergence during contraction phases and to secondary contact(s) and genetic admixture during expansion phases (Nevado et al. 2018; Wielstra, Salvi, and Canestrelli 2021; Zemlak et al. 2008). This played a major role in shaping the distribution of genetic variation of closely related boreal species. Paleoecological and genetic studies have shown that the impact of glacial cycles varied extensively across species and geographical areas. In western Europe, for instance, some species could only survive in the Iberian or Italian peninsulas while others were able to survive as far north as Central Europe (Binney et al. 2009; Willis, Rudner, and Sümegi 2000). The extent of glaciation also varied through space: while Fennoscandia was almost fully glaciated during the Last Glacial Maximum (LGM, 26, 500-19,000 years ago), Siberia was not. Siberia was a cold desert where pockets of trees were able to survive (Semerikov et al. 2013 and reference therein). These repeated sequences of reproductive isolation and admixture, and the vast but non-homogeneous population movements that accompanied them at continental scales, have led to the accumulation over time of an ancient, deep-seated and complex genetic structure, as, for instance, observed in seven European tree species (Milesi et al. 2023). This structure was blurred by extensive gene flow, or reinforced by strong local adaptation, both of which are characteristics of most forest tree species (Savolainen, Pyhäjärvi, and Knürr 2007). This already multi-layered structure has often been further enhanced by hybridization.

Over the last decades, extensive genome-wide resequencing studies have highlighted the extent of hybridization and introgression between closely related species (Belokon et al. 2022; N. Chen et al. 2018; Cullingham et al. 2012; Fu et al. 2022; Keim et al. 1989; Sankararaman et al. 2014; Shuvaev et al. 2023; Thórsson, Salmela, and Anamthawat-Jónsson 2001). Secondary contacts -when previously isolated populations are re-united-have been extensively studied, in particular for their role in speciation, for instance through reinforcement of reproduction isolation (e.g. (Barton and Hewitt 1985)). When not selected against, species hybridization is expected to increase genetic diversity and can allow for the transfer of beneficial genetic material from one genetic background to another (Chhatre et al. 2018; Jones et al. 2018; Leroy et al. 2020; Platt et al. 2019; Whitney, Randell, and Rieseberg 2010) directly affecting phenotypic traits (e.g., Darwin’s finches beak shape, Grant and Grant 2019) and thereby allowing for niche expansion (e.g., (Pfennig, Kelly, and Pierce 2016) and reference therein). In turn, ecological niche expansion, could result in an increase in census population sizes and foster populations resilience to global changes. In order to better understand the interplay between hybridization and demography history one has thus to investigate simultaneously the evolutionary dynamics of the various genetic entities/species involved. Doing so require extensive sampling across entire distribution ranges and examples of such large-scale study remain scarce outside model species (e.g. (Bruxaux et al. 2023)).

In the present study we combined different methods to reconstruct the joint demographic history of two conifer species, Norway spruce (*Picea abies* [L.] H. Karst.) and Siberian spruce (*P. obovata* Ledeb). Norway spruce and Siberian spruce are keystone species of the Eurasian boreal forest, and their joint range extends from the Norwegian Coast in the West to the Sea of Okhotsk in the East. Similar to other boreal tree species, Norway and Siberian spruce experienced cycles of population expansion and contraction over multiple glacial periods. The current distribution of Norway spruce and Siberian spruce shows a clear West to East pattern but no clear delineation between their natural range can be drawn; they occupy different but overlapping ecological niche (Karunarathne et al. 2023). Phenotypic traits also show a continuous distribution from one genetic background to the other (e.g. survival, growth, shape of cone-scale, Lagercrantz and Ryman 1990; E. Nakvasina et al. 2019; Orlova et al. 2020; Popov 2010) and Norway spruce can be discriminated from Siberian spruce and from their hybrid forms (E. Nakvasina et al. 2019; E. N. Nakvasina, Volkov, and Prozherina 2017; Popov 2010, 2013). Still, in spite of their paramount economic and ecological importance, the geographic patterns of genetic variation of the two species remains unclear, in particular in their putative contact zone in the western Russian plains.

While *P. abies* can be separated into several well delineated genetic clusters reflecting population movement during glacial cycles and main geographic domains (J. Chen et al. 2019; Tsuda et al. 2016), no clear population subdivision was so far observed within *P. obovata* using nuclear markers despite its much wider range (Tollefsrud et al. 2008, 2015; Tsuda et al. 2016). The weaker population structure in *P. obovata* could reflect more stable populations during ice cycles, but it could also be explained by restricted geographical sampling and genomic surveys. Current literature suggests that the two species entered into contact after LGM with a strong East to West recolonization of *P. obovata*, introgressing *P. abies* populations in their northern range (J. Chen et al. 2019; Li et al. 2022; Tsuda et al. 2016). By jointly analyzing nuclear (microsatellites) and mitochondrial markers, Tsuda et al. (2016) evidenced two migration barriers, one roughly separating the northern from the southern range of Norway spruce and the other corresponding to the Ural Mountains. These two geographic barriers were suggested to be responsible for the observed non-homogeneous longitudinal gene flow between the two species. The geographic discrepancy between nuclear and mitochondrial markers also suggested more complicated evolutionary history than a single recent secondary contact between two previously isolated genetic background during LGM. The mitochondrial haplotypes of the northern domain of *P. abies* group together with the ones from *P. obovata* and are clearly separated from *P. abies* haplotypes from the southern domain. However, in contrast, nuclear markers clearly separate *P. obovata*, on one hand, from *P. abies* Northern and Southern domains, on the other hand (Tollefsrud et al. 2008, 2015). As mitochondria are maternally inherited, *i.e.* are dispersed only through seeds, it is unlikely that the northern admixture between *P. abies* and *P. obovata* only reflects a recent contact between the two species. Extensive genomic data confirmed the presence of gene flow from *P. obovata* into the northern domain of *P. abies* and suggested the existence of four main genetic ancestral clusters across the *P. abies* -*P. obovata* complex (J. Chen et al. 2019; Li et al. 2022). Unfortunately, a highly biased sampling towards *P. abies* populations and a limited sampling of individuals located within the putative hybrid zone strongly limited inferences on species genetic interaction over multiple ice-cycles.

Here we analyzed the genome-wide variation (480,428 nuclear SNPs and 87 chloroplast SNPs) of 542 individuals sampled in 55 populations, ranging from Western Europe and Fennoscandia to the Yenisei River in Siberia, with a particularly dense sampling of the putative *P. abies*-*P. obovata* hybrid zone. We first describe the current spatial variation in genetic diversity across both species range and revealed a strong population structure in *P. obovata* also, notably along the Ural Mountains. This structure differs markedly from that of *P. abies*. We then characterized the hybrid zone between the two species and showed that it extends from Fennoscandia in the northwest all the way to southern Ural Mountains in the southeast. Estimation of isolation by distance and migration surfaces showed biased introgression of *P. obovata* into the northern domain of *P. abies* and of *P. abies* into the southern domain of *P. obovata.* This pattern is supported by long range gene flow carried over by pollen dispersal as suggested by the analysis of chloroplast DNA markers. Finally, a joint demographic analysis of the two species supported the presence of multiple contacts over glacial cycles and synchronous changes in population dynamics, giving rise to new genetic clusters that persisted until today.

## Material and Methods

### Sampling, sequencing and SNP calling

In this study, a total of 542 trees from 55 populations were sampled across *P. abies* and *P. obovata* natural ranges (from 41.8° to 69.0°North and from 8.2° to 92.8°East), with a particularly dense sampling within the putative large hybrid zone between the two species (Fig. S1). Needles of each sample were collected and used for DNA extraction using the DNeasy Plant Mini Kit (QIAGEN). Genomic DNA was then submitted to exome capture with 40,018 probes, targeting 26,219 genes (Vidalis et al. 2018). Raw-reads are available in SRA (https://www.ncbi.nlm.nih.gov/sra) under bioproject number PRJNA511374 and PRJNA1007582. Paired-end Illumina libraries were constructed and sequenced by RAPiD Genomic, USA. Clean reads were aligned to the *P. abies* reference genome v1.0 (Nystedt et al. 2013) using BWA with default parameters. PCR duplicates were removed using PICARD v 2.27.4 (https://github.com/broadinstitute/picard/), followed by genotype identification carried out with GATK HaplotypeCaller individually and SNP calling using GenotypeGVCFs across all samples jointly. Hard-filtering was performed to filter out SNPs of low quality, based on the following criteria: QD < 2.0, MQ < 40.0, SOR > 3.0, QUAL < 20.0, MQRankSum < -12.5, ReadPosRankSum < -8.0, and FS > 60.0. Only putatively neutral SNPs (i.e. SNPs located in intron or intergenic regions, or being synonymous in protein coding sequences) with call rate larger than 0.7 and minor allele frequencies larger than 0.01 were kept for further population structure and demographic inference analysis. After filtering steps, 1,761,609 SNPs were retained across 542 spruce individuals. We then applied further filtering steps to keep only unlinked putatively neutral SNPs (*r*^2^ < 0.5, window size of 50 SNPs) using Plink v2.0 (Purcell et al. 2007). The final dataset contains a total of 480,428 putatively neutral SNPs. Polymorphic sites located within the paternally inherited chloroplast sequence were also identified following the same protocol save for pruning for high linkage disequilibrium. Only 484 samples were kept for the chloroplast dataset while the rest were filtered out due to high missing rates (>30%).

### Population genetic structure and clustering

Population genetic structure was first investigated using the clustering algorithms implemented in *ADMIXTURE* v. 1.3.0 (Alexander, Novembre, and Lange 2009) based on putatively neutral SNPs. Different numbers of ancestral components (K) were tested ranging from one to ten. We performed ten-fold cross validation and 200 bootstraps to define the theoretical best K value and selected the one with the lowest cross validation error. As ADMIXTURE/STRUCTURE method was revealed to be less appropriate for continuously distributed species with isolation by distance (Novembre and Stephens 2008), which is the case for *P. abies* and *P. obovata*, we also ran a Principal Component Analysis (PCA) to assess spatial genetic patterns using EIGENSOFT v. 7.2.0 with default parameters (Galinsky et al. 2016). Fine-scale genetic clustering was then performed with Uniform Manifold Approximation and Projection (UMAP) analysis (Diaz-Papkovich et al. 2019). Briefly, UMAP groups genetically similar individuals together on a local scale by creating a neighborhood around each individual’s coordinates and also preserves long-range topological connections to more distantly related individuals. Therefore, the number of neighbors and minimum genetic distance could lead to different clustering relations. The top five principal components (PCs) were retained in UMAP analysis. Default values were used for dimensions (NC=2). We tested different minimum genetic distance (MD, 0.001, 0.01 and 0.1) and neighbor numbers (NN, 5, 10, 15). All combinations produced comparable results and we retained the result with MD = 0.1 and NN=10 as this combination showed the most discriminant clustering.

In order to compare with genetic clustering when populations, instead of individuals as above, are considered, we used Bayesian inference implemented in the BayPass software (v2.2 – BayPass Gautier 2015) to estimate the empirical patterns of covariance in allele frequencies between the populations (see also Günther and Coop 2013). To minimize the effect of linkage disequilibrium, we divided putatively neutral SNPs into subsets of 20,000 randomly selected SNPs and the covariance matrices were generated using 50,000 MCMC iterations for all subsets. For each subset independently, the last 1000 matrices after burn in were averaged, followed by averaging all the sub-matrices across subset.

### Genetic diversity and population divergence estimation

For each population (*i.e.* sampling location) and each genetic cluster identified using *UMAP* approaches we computed nucleotide diversity (Nei’s π, Nei and Tajima 1981) and Tajima’s D (Tajima 1989) using the R package *‘PopGenome’* v2.7.5 (Pfeifer et al. 2014). We also estimated Weir and Cockerham weighted *F_ST_* (Weir and Cockerham 1984) for each population pair using *VCFtools* v0.1.17 (Danecek et al. 2011).

### Isolation-by-distance and effective migration surfaces

To further explore the dispersal/introgression pattern of the two species within the sampling area, we investigated the pattern of Isolation-by-Distance (IBD, Morton 2013). Population pairwise geodesic distances were calculated using R package ‘*geodist*’v0.08 (Padgham 2021).

The global pattern of genetic dissimilarity decay across space was estimated by regressing *F_ST_* /(1-*F_ST_*) for each population pair as a function of the geodesic distance between the corresponding populations. We also ran a Mantel test (Mantel 1967) with 10,000 permutations to test for the significance of the correlation between the genetic and geodesic distance matrices using R package ‘*vegan’* v2.6 (Dixon 2003).

Perturbation to gene flow (i.e. barriers and corridors to gene flow) might lead to deviation from the expectation of IBD. We thus identified corridor and barrier to gene flow using the *fEEMS* v1.0.1 software (Marcus et al. 2021) that allows estimating effective migration surfaces across space. We first generated an outer coordinate file of the sampling area using the polyline method in the google maps api v3 tool (http://www.birdtheme.org/useful/v3tool.html). We adopted a grid size of 100 km^2^ that was the best compromise between computational burden and model mis-specification. All 55 populations were positioned into 48 different grids. Populations within the same grid were genetically closely related, leaving minor risk of bias in migration surface estimation with adjacent grids. Cross validation was performed and the number of iterations automatically stopped once convergence was achieved. The tuning parameter lambda controls the strength of penalization placed on the output of the migration surface. We chose the value of lambda that minimizes the cross-validation error.

### Chloroplast haplotype group analysis

In *P. abies* and *P. obovata*, chloroplasts are paternally inherited through pollen. Investigating the distribution of genetic variation of chloroplast sequences thus provides additional information on pollen dispersal and long-distance gene flow (Scotti et al. 2008; Tollefsrud et al. 2015). SNPs located in chloroplast genome were used for unrooted maximum likelihood (ML) phylogenetic inference using *IQtree* v2.03 (Nguyen et al. 2015). Node support was measured with 1000 replicates of ultrafast bootstrap and 1000 replicates of Shimodaira-Hasegawa-like aLRT (SH-aLRT) test (Guindon et al. 2010).

### Inference of ancient migration events between the main clusters

We used *TreeMix* v1.13 (Pickrell and Pritchard 2012) to infer the pattern of population splits and major historical migration events among the main genetic entities. A maximum likelihood phylogenetic tree of the main genetic clusters identified using *UMAP* was first built by *TreeMix*. The support of the resulting phylogeny was evaluated by bootstrapping blocks of 500 SNPs. The homogenous genetic cluster formed by populations located along the Yenisei River was used as the outgroup to root the tree. Up to ten edges (migration events, parameter “*m*” in *TreeMix*) were then added to the dendrogram when two populations are more closely related to each other than expected under a model of bifurcating splitting. Branches were then rearranged after each new edge was added. Ten bootstrap replicates were performed for each number of edges. The optimal number of admixture events was determined when the model explained over 99.8% of the variance to avoid model overfitting. The inferred migration events were verified using f3-tests implemented in *ADMIXTOOLS2* (Patterson et al. 2012).

### Demographic history inference of the main clusters

*TreeMix* was used to infer reticulate evolution on the basis of a maximum likelihood phylogenetic tree which provided minor information about the timing of the admixture event. A proper outgroup (*e.g.* a population from a third species) was also missing, making it difficult to interpret the relationship between groups. In order to test alternative demographic models, we used the coalescent-based composite likelihood method implemented in *FastSimCoal2* (Excoffier et al. 2021). When estimating the parameters of different models, we used a generation time of 25 years and a mutation rate of 2.75 x 10^-8^ per site per generation (Ann et al. 2007; J. Chen et al. 2012, 2019; Nystedt et al. 2013).

We defined the models to be tested based on the output of the *TreeMix* analysis. The models were built with increasing complexity and migration events were added progressively in the order suggested by the *TreeMix* analysis. Five scenarios with different number of edges added (ranging from 0 to 4) were initially tested. We then tested additional models, notably considering different origins for the clusters with ambiguous phylogenetic positions (e.g, admixture *vs* split). Migration was only allowed between adjacent clusters to avoid over-parametrization (see Supplementary material 1 for model comparison).

*ANGSD* v0.933 (Korneliussen, Albrechtsen, and Nielsen 2014) was used to estimate pairwise 2-dimensional SFS as input to *FastSimCoal2*. Estimates of the parameters of the various demographic models were optimized with Brent’s algorithm with 50 iterations and 100,000 simulations for each iteration. The models’ goodness of fit was then evaluated using the likelihood ratio G-statistics (CLR = log_10_(CL_O_/CL_E_), where CL_O_ and CL_E_ are the observed and estimated maximum composite likelihood, respectively (Excoffier et al. 2013). To estimate 95% confidence intervals, we first used a large range of values for parameters prior and investigated their posterior distribution after 200 bootstrap replicates. We then restricted the range of parameter prior for another set of 200 bootstrap replicates and computed the 95% confidence interval from parameters posterior distributions.

We then investigated whether changes in effective population size (*Ne*) were synchronous between the two species, as would be expected under glacial cycles, by inferring the dynamics of *Ne* changes over generations using *StairwayPlot2* (Liu and Fu 2020). Folded SFS for 12 selected populations from the most divergent genetic clusters (8 populations from EUR cluster: DE_1, DE_2, CZE, GERH, GERL, SK, RO_1, RO_2; 4 populations from YEN cluster: BOR, TYR, KRA, ENI) were estimated with *realSFS* implemented in *ANGSD* v0.933. Having no reason to consider different generation times or mutation rates between *P. abies* and *P. obovata* we directly compared the raw output from *StairwayPlot2* (i.e. assuming a generation time of *Gt* = 1).

## Results

### Well defined genetic clusters despite limited genetic differentiation

We investigated the population structure by first performing a PCA analysis on genomic variations of all samples (N=542 with 480,428 putatively neutral nuclear SNPs, Fig. S2). The first two PCs together explained only ∼5.1 % of the total genetic variation but genetic clustering reflects geography. PC1 mainly discriminates populations along a longitudinal gradient (Spearman’s *rho*=-0.94, S= 51 150 760, *p* < 0.001) with populations from the Alpine domain (*P. abies*) at one extreme and populations from the Yenisei River (*P. obovata*) at the other. Change from one genetic cluster to another is rather continuous and no clear delineation between the two species can be drawn. PC2 shows lower but strongly significant correlation with latitude (Spearman’s *rho*= 0.35, S= 17 045 820, *p* < 0.001), in particular within *P. abies* and hybrids populations. As some population structure is still captured by other principal components (*e.g.* PC3 and 4 see Fig. S2), we performed dimensionality reduction incorporating the first five PCs using *UMAP*. Nine genetic clusters can be distinguished among which two include only one or two populations (Fig. 1 and Supplementary material 2). The first one (ITA) contain two Italian populations (ITAL and ITAH) isolated from the rest of the *P. abies* main range (Fig. 1 and S1). The second one (PAH) is made up of individuals coming from an elevated plateau in Southern Urals (PAH population on Mount Iremel). These two clusters are isolated from the rest of the range and experienced stronger genetic drift. The seven other clusters globally grouped populations according to their geographical distribution and were named after the main geographical domains (Fig. 1). The Yenisei (YEN) cluster only contain individuals from populations along the Yenisei River and stands out by its genetic homogeneity despite the large latitudinal distribution of the populations (∼10 latitudinal degrees). In comparison, populations from Urals Mountains and Ob River show much more structure for a similar latitudinal range and URAL and OB clusters mainly separate Southern populations from Northern ones. Similarly, the main population structure appears along latitude for *P. abies* populations. EUR cluster gathers Southern Europe populations from the Alpine and Carpathian ranges, and EUR_RUS cluster groups populations from the Russia-Baltic region. Fennoscandia domain is split into a Northern (Fenno1) and a Southern (Fenno2) genetic clusters that expands up to the Western plains of Russia and Southern Urals, respectively.

**Figure 1.**
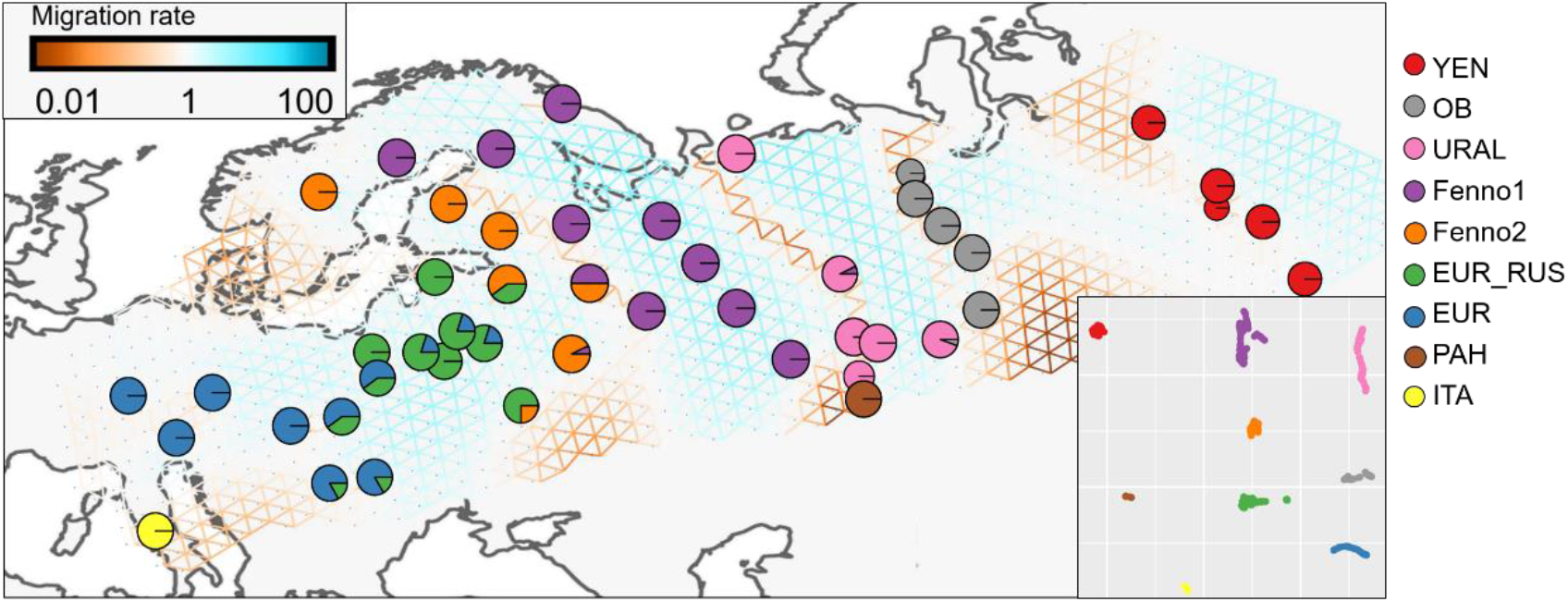
Population structure and gene flow. Geographic distribution of the nine genetic clusters and estimated effective migration surface across space. Genetic clusters definition was based on the results of *UMAP* shown as inset.

We then adopted a population-based clustering approach (Günther and Coop 2013). Bayesian inference revealed a similar geographic pattern of genetic clustering as *UMAP*’s. Two main groups were identified, corresponding to populations with more *P. abies* background (PA) or *P. obovata* background (PO), respectively, and nine well delineated clusters were identified (Fig. S3). The main difference compared with the clustering made at individual level is that two populations only containing individuals from URAL and two only from EUR_RUS cluster separately from the rest and that ITA and PAH cluster now cluster with geographically close by populations. Based on individual clustering (PCA - UMAP) most populations gather individuals originating from a single cluster, but ten populations from Central and Eastern Europe (e.g. populations from PA:2.1.2 and PA:2.1.3 in Fig. S3) contain individuals from both EUR and EUR_RUS clusters or from both EUR_RUS and Fenno2 clusters (Fig. 1). This suggests either recent secondary contacts through post-LGM population expansion or genetic transfer linked to human activities. As it might affect downstream analyses, for instance inference of demographic history, we decided to use the genetic cluster defined at the individual level (PCA-UMAP).

In order to better understand the origin of this population structure, we ran an *ADMIXTURE* analysis to gain further insights into the ancestry components of today’s clusters. If admixed populations were simply the result of recent or ongoing hybridization and introgression between *P. abies* and *P. obovata* genetic backgrounds, we can expect a best theoretical number of K = 2 ancestry components. Based on five-fold cross-validation (CV), we found that K = 3 or 4 best explained the data (Fig. 2B). For both K= 3 and 4, *ADMIXTURE* revealed a continuous genetic composition change from “pure” EUR to “pure” YEN genetic background, which is consistent with the result of the PCA analysis. However, in contrast with what would be expected if K = 2, the admixed populations do not directly share EUR and YEN genetic components. Instead, the analysis revealed a two-fold admixture pattern, with populations sharing either EUR (EUR_RUS, Fenno1 and 2) or YEN (PAH, URAL and OB) genetic component with a cluster of third origin (Fig. 2A). The main ancestry component in the hybrid zone for K = 3 (green, Fig. 2A, top panel) split into two components, AC_UR and AC_FEN, for K=4 (Fig. 2A, bottom panel). In any case, *ADMIXTURE* analysis suggests that at least one ancestry (K=3) or two closely related ancestries (K=4) found in the current hybrid zone played an important role in shaping the distribution of genetic variation of the two-spruce species and that the evolutionary dynamics of the two species are probably more intricate than simply a secondary contact after LGM.

**Figure 2.**
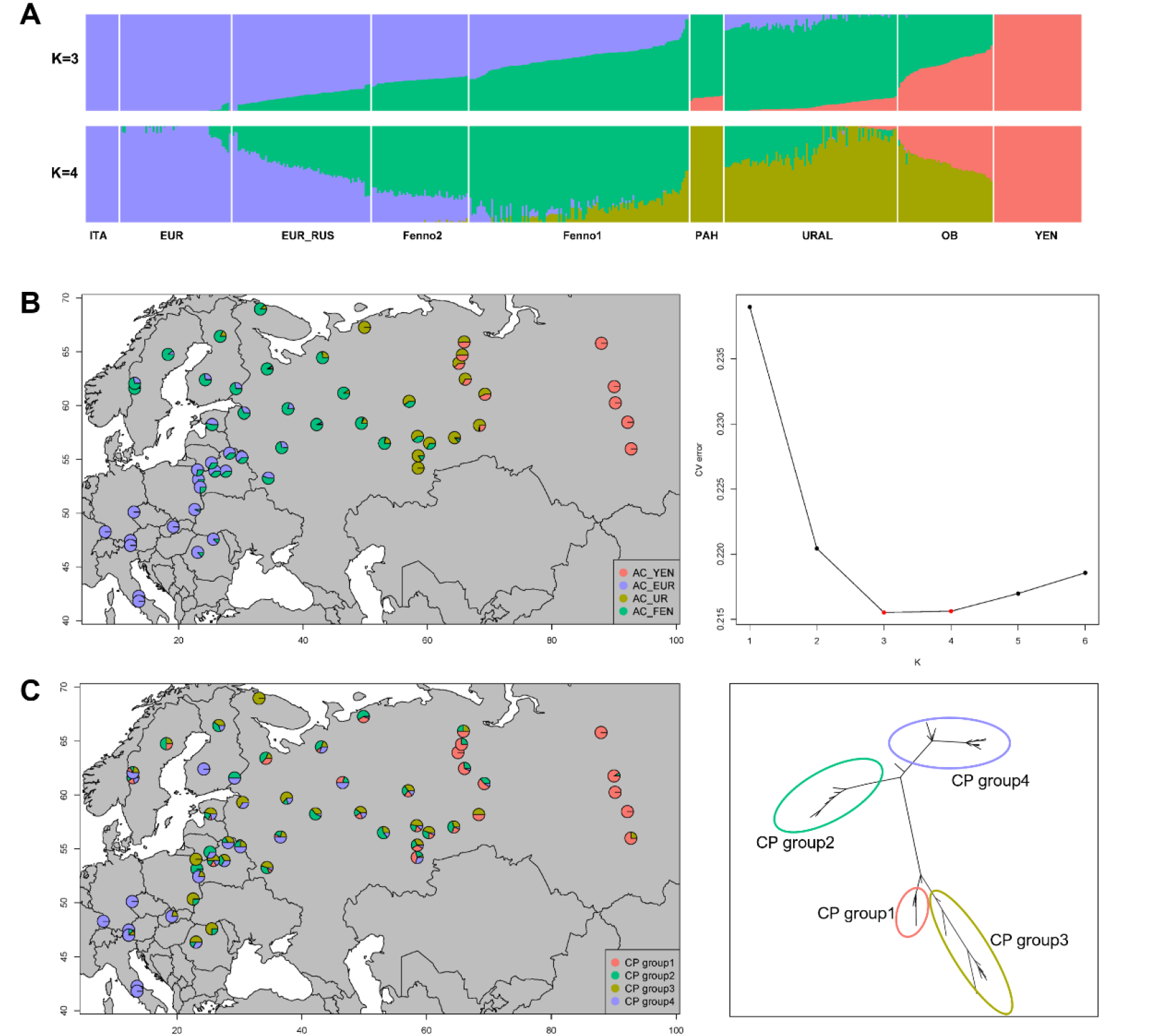
Distribution of ancestry component and chloroplast haplotypes. **A.** *ADMIXTURE* plot showing the ancestry components (nuclear DNA) distribution for K = 3 and K = 4. **B**. Geographic distribution of ancestry components (left) and scree plot of cross-validation errors for different K values in *ADMIXTURE* analysis (right). **C.** Geographic distribution of the four main chloroplast DNA haplotype groups (left) and maximum likelihood tree showing the relationship among the four main haplotype groups (right).

### Distribution of genetic diversity

We estimated Nei’s π and Tajima’s D at both population and genetic cluster levels. At population level, the estimated π ranges from 0.00524 to 0.00687, with a median of 0.00624, and estimates of π were globally lower for *P. abies* than *P. obovata* populations (Supplementary material 3). The nucleotide diversities increased from West to East (Spearman’s *rho*=0.5, S=13 413, *p* < 0.001) with the populations from the main hybrid zone (Western Russia populations) exhibiting an intermediate level between main European *P. abies* populations and Yenisei River *P. obovata* populations (Supplementary material 3); π also increased with latitude (Spearman’s *rho*=0.1, S=19 165, *p* = 0.02), but the relationship is pulled by *P. abies* alone. The same relationship as that at population level is observed at the cluster level and, considering their geographical isolation, unsurprisingly, the ITA cluster exhibited the lowest nucleotide diversity (0.00589) among pure “*P. abies*” clusters and PAH the lowest π (0.00627) among pure “*P. obovata*” clusters (i.e. YEN, PAH, URAL and OB) (Tab. 1).

**Table 1.** Proportion of ancestry components and summary statistics for the nine genetic clusters.

| clusters | AC_UR | AC_EUR | AC_FEN | AC_YEN | Tajima's D | pi ($10^{-3}$) |
| --- | --- | --- | --- | --- | --- | --- |
| ITAH | 0 | 1 | 0 | 0 | -0.471 | 5.89 |
| EUR | 0 | 0.96 | 0.04 | 0 | -1.092 | 6.14 |
| EUR_RUS | 0 | 0.57 | 0.43 | 0 | -1.158 | 6.3 |
| Fenno2 | 0 | 0.29 | 0.71 | 0 | -1.096 | 6.46 |
| Fenno1 | 0.12 | 0.03 | 0.85 | 0 | -1.04 | 6.36 |
| PAH | 1 | 0 | 0 | 0 | -0.415 | 6.27 |
| URAL | 0.77 | 0 | 0.22 | 0.01 | -0.981 | 6.42 |
| OB | 0.61 | 0 | 0 | 0.39 | -0.765 | 6.7 |
| YEN | 0 | 0 | 0 | 1 | -0.213 | 6.56 |

Tajima’s D values are negative in all populations (-0.91 – -0.031, median: -0.397) but not significantly different from 0 considering that Tajima’s D roughly follows a beta distribution of mean 0 and variance 1 (Tajima 1989). This suggests that the populations do not significantly depart from the standard coalescent model, although the global negative trends are consistent with recent population expansions after the LGM. This is confirmed at the cluster level where *P. abies* clusters exhibit lower Tajima’s D than *P. obovata* clusters. The highest Tajima’s D is observed for the YEN cluster, probably because its populations were less affected by glacial cycles than Western European ones (Tab. 1).

### Extensive but non-homogeneous gene flow across space

Our population structure/admixture analyses suggest both extensive ancient admixture events and ongoing gene flow between *P. abies* and *P. obovata* genetic backgrounds. To tease apart the effect of each on the current genetic diversity distribution we first investigated the pattern of Isolation-by-Distance (IBD) by regressing pairwise *F_ST_* / (1 - *F_ST_*) over the geodesic distance between each population in a pair. The divergence among populations was generally low with a highest *F_ST_* close to 0.25 found between Yenisei River and Italian populations. The regression revealed a striking IBD pattern across the whole range of both species (Fig. S4, Mantel’s correlation coefficient *r*: 0.92, *p* < 0.001), and no gap in the distribution was observed as might have been expected if hybridization between *P. abies* and *P. obovata* was limited. We then used the linear regression between *F_ST_* / (1-*F_ST_*) and geodesic distance (*r*^2^ = 0.85) as a baseline to look for area where gene-flow is stronger (below the baseline) or weaker (above the baseline) than the global trend across the whole distribution range (Fig. S4). Two main deviations where identified. Populations located in Fennoscandia and those located in Europe mainland (Alpine and Carpathian ranges, Poland and Baltic states, see Fig. S1) show a closer relationship than expected (*t*-test between the global residual and local residual, *t* = -11.896, *df* = 296.46, *p* < 1e-5) while Western Russian populations and Yenisei River populations are more distant than expected from the global IBD pattern (*t* = 11.068, *df* = 66.32, *p* < 1e-5).

To further identify barriers and corridors to migration, we used *fEEMS* to estimate the geographical distribution of effective migration rate (Fig. 1). As expected, areas with reduced migration rate (thereafter called barriers) delineate the main genetic clusters identified with *UMAP*, while the migration rate was higher than average within clusters. Within *P. obovata*, the Ural Mountains clearly constitute a barrier to gene flow separating URAL and OB genetic clusters. In *P. abies*, Fenno2 is distributed around the Baltic Sea that was identified as a barrier to gene flow. However, no clear topological barrier can be found to explain the separation of EUR_RUS from EUR. Importantly, we found two main barriers to gene flow delineating a corridor between Southern Urals and Northern Fennoscandia that do not correspond to any clear topological variation. The first barrier separates Fenno2 from Fenno1 in the West and the second separates Fenno1 from URAL in the East (Fig. 1). These two barriers are probably explained by a combination of ancient demographic events and ecological transition, for instance, the barrier in the west matches the Yekaterinburg-Murmansk climatic transition.

### Distinct geographic distribution patterns between genomic ancestries and main chloroplast groups

Four main well-supported chlorotype groups were identified from the plastomic ML phylogeny (Fig. 2C). Two of them represent the main chlorotypes from *P. obovata* (CP group1 and 2), the two others belong to *P. abies* (CP group3 and 4). The geographic distribution of the chlorotype groups globally matches the distribution of the ancestry components (admixture analysis, K = 4) but with a larger spread. Considering that chloroplast is paternally inherited in the two spruce species, the observed pattern thus confirms the potential for long-range wind dispersal of pollen. For instance, CP group4 and CP group1, that are predominantly found in Western Europe and Yenisei River populations, respectively, are also found in populations located within the hybrid zone while the two corresponding main ancestry components estimated from nuclear DNA, AC_EUR and AC_YEN, are not into contact in the hybrid zone. The other two groups (CP Group2 and 3) are widely distributed across the hybrid zone with no clear geographic pattern despite the fact that they belong to the two different species (Fig. 2C), in contrast with nuclear DNA that shows much more structure in the hybrid zone (see AC_FEN and AC_UR, Fig. 2B). Finally, most populations from the hybrid zone display chlorotypes belonging to at least three different groups. *P. obovata* chlorotypes introgressed further West in the North and those from *P. abies* further East in the South, contributing to the active bi-directional, but asymmetric, gene flow occurring in the hybrid zone.

### **Biased ancient migration from *P. obovata* into *P. abies***

The previous analyses suggested the occurrence of ancient admixture events between *P. abies* and *P. obovata.* We used *TreeMix* to quantify the number, intensity and direction of main gene flow events between the two species. The best model explained 99.8% of the total variance and added four edges (i.e. main migration events) to the initial tree (Fig. 3A and S5). The resulting phylogenetic tree first separates YEN and OB from EUR, Fenno1, Fenno2, and EUR_RUS clusters with URAL branching out from the branch connecting these two groups. EUR_RUS cluster splits early from the common ancestor of EUR, Fenno1 and Fenno2 clusters. The model suggests that the first migration event occurred from *P. obovata* toward what is today the northern range of *P. abies* (Fenno2, arrow 1 in Fig. 3A). More importantly, we detected two concomitant migration events occurring from URAL towards OB (*P. obovata*) and Fenno1 (*P. abies*) (arrows 2 and 3, respectively, in Fig. 3A). URAL is thus an ancestral cluster that influenced both *P. obovata* and *P. abies* genetic diversity, shedding a new light on the patterns observed with *ADMIXTURE* and *fEEMS* analyses. The most recent migration event was from EUR towards EUR_RUS (arrow 4, Fig. 3A). This explains the close relationship between the two clusters observed with PCA and *ADMIXTURE*, despite the long divergence time suggested by *TreeMix*. All migration events were confirmed with f3 test as suggested by (N. Chen et al. 2018) (Supplementary material 4). However, one needs to be cautious when interpreting the reticulate evolutionary history revealed by *TreeMix*. In particular, when the migration weights are close to 0.5, the branching and the source population for migration could be flipped, both being theoretically indistinguishable, see for instance arrows 3 and 4 in Fig. 3A.

**Figure 3.**
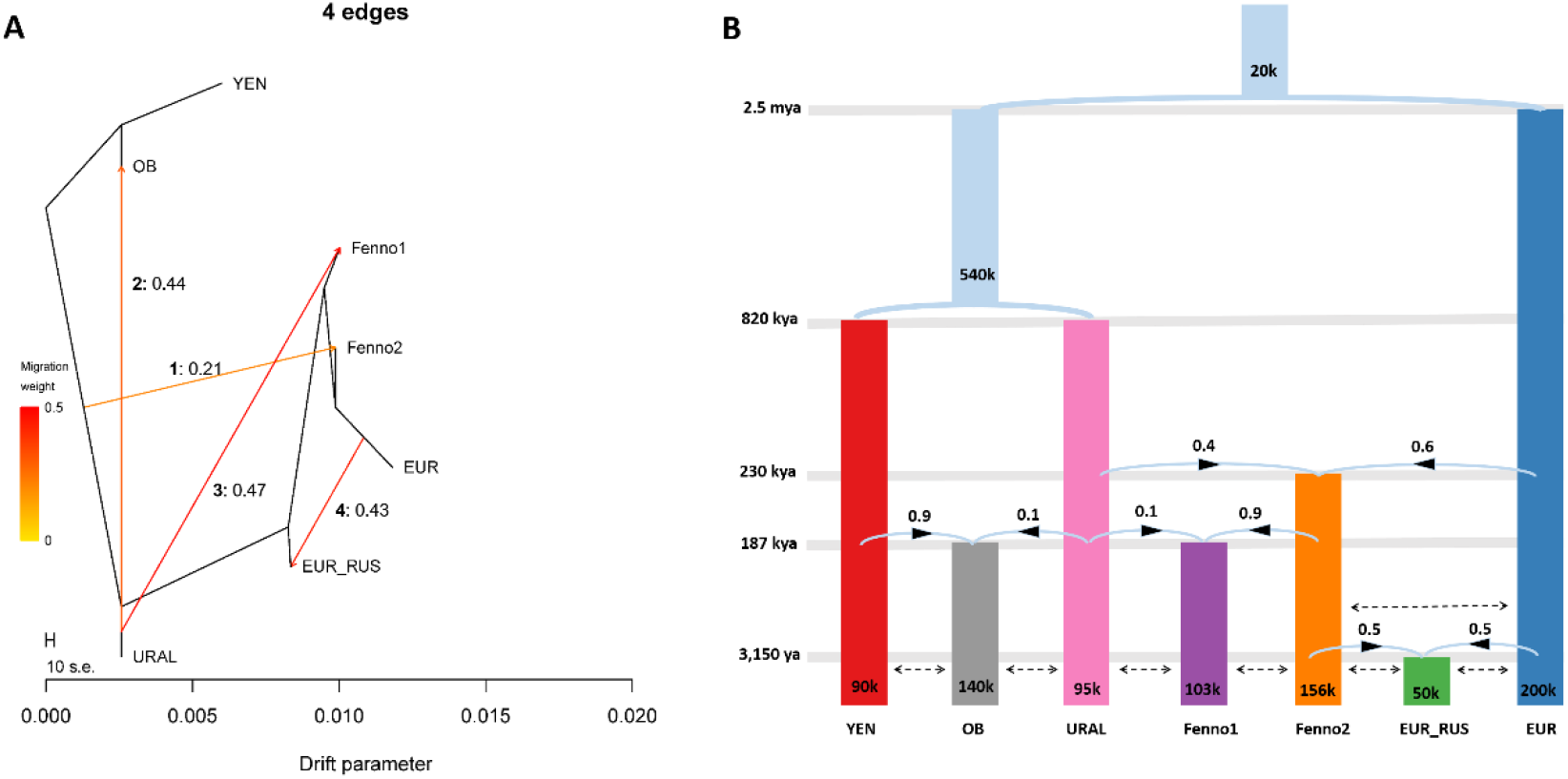
Inference of main demography history. **A.** *TreeMix* analysis, the tree with the highest support considers four main migrations event; **B.** *FastSimCoal2* analysis, the best model with four admixture events is presented. For both plots, the numbers above/next to arrows indicate the relative contribution of the source populations to the sink populations.

### Demographic history inference

The results of *TreeMix* provide new insights into the complex evolutionary dynamics of the two species but did not allow for a detailed inspection of their demographic histories. In particular, it is not clear whether UR1 split from YEN or EUR. Similarly, the origin of EUR_RUS remains unclear due to the roughly equal contribution of source populations. To solve these issues, we explored different demographic scenarios with different sets of parameters and topologies with *FastSimCoal2* (supplementary material 1). Among the 36 models we explored, one model stands out and recovered the observed site frequency spectra well (CLR: -0.0027, Supplementary material 5). The best model and the confidence intervals for the estimated parameters are presented in Fig. 3B and Tab. 2.

**Table 2.** Demographic parameters estimated for the model using *fastsimcoal*. N, effective population size; M, migration proportion from source cluster to sink cluster for admixture events; T, time for divergence or admixture events. Tanc, coalescent time for all clusters.

| Parameters | Point estimation | 2.50% | 97.50% |
| --- | --- | --- | --- |
| N<sub>EUR</sub> | 200, 301 | 198, 875 | 268, 481 |
| N<sub>OB</sub> | 139, 799 | 123, 490 | 169, 277 |
| N<sub>YEN</sub> | 90, 266 | 63, 134 | 148, 423 |
| N<sub>URAL</sub> | 95, 476 | 91, 911 | 142, 727 |
| N<sub>Fenno1</sub> | 103, 151 | 97, 447 | 112, 885 |
| N<sub>Fenno2</sub> | 156, 520 | 152, 277 | 202, 673 |
| N<sub>EUR_RUS</sub> | 50, 108 | 50, 386 | 123, 478 |
| M<sub>YEN-OB</sub> | 0.94 | 0.81 | 0.98 |
| M<sub>EUR-Fenno2</sub> | 0.60 | 0.56 | 0.69 |
| M<sub>URAL-Fenno1</sub> | 0.08 | 0.02 | 0.18 |
| M<sub>EUR-EUR_RUS</sub> | 0.46 | 0.46 | 0.49 |
| T<sub>Fenno2</sub> | 230, 650 | 214, 191 | 281, 793 |
| T<sub>URAL</sub> | 818, 900 | 571, 852 | 1, 088, 353 |
| T<sub>Fenno1</sub> | 188, 650 | 181, 676 | 206, 930 |
| T<sub>OB</sub> | 187, 225 | 105, 660 | 244, 877 |
| T<sub>anc</sub> | 2, 439, 200 | 2, 300, 048 | 2, 790, 453 |
| T<sub>EUR_RUS</sub> | 3150 | 3, 411 | 9, 683 |

The use of a coalescent-based approach confirmed that the UR1 cluster is ancient (T_UR1_: ∼820 kya) and diverged from the YEN cluster after the split between the ancestors of Norway and Siberian spruces (T_anc_: ∼2.5 mya). It also emphasizes the importance of admixture events in the history of the two species. Hence, both Fennoscandia clusters are of hybrid origin. Fenno2 resulted from the admixture between URAL and EUR (T_Fenno2_: ∼230 kya) with a higher contribution from the *P. abies* genetic background (relative contribution, 60%) while Fenno1 originated slightly later from the admixture between URAL and Fenno2 (∼187 kya), with a major contribution from Fenno2 (∼90%). At roughly the same time, the OB cluster formed from the admixture between YEN (∼90%) and URAL (∼10%). The concomitance of these two events suggests that both events occurred during the same interglacial period. Finally, the EUR_RUS cluster resulted recently from the admixture between EUR (∼50%) and Fenno2 (∼50%) clusters, likely as a result of the post-LGM population expansion (T_EUR_RUS_: ∼3,150 years ago).

In summary, both *TreeMix* and *FastSimCoal2* analyses show consistent results and highlight three key features in *P. abies* and *P. obovata* evolutionary history. First, most of the clusters originated much before the LGM. Second, the main clusters were the results of admixture between more ancient genetic entities, probably as a consequence of the Pleistocene ice-cycles. Third, recurring introgression between the two spruce species played a critical role in shaping the current distribution of genetic variation and with a central role played by the URAL cluster that contributed to several admixture events in both species.

As indicated above, the parameters estimated with *FastSimCoal2* suggested multiple admixture and introgression events in the evolutionary history of Norway and Siberian spruce, some happening at the same time and thus likely following glacial periods. Still, it is not clear if the admixture events were due to the movement of a single genetic entities toward the other or if both species experienced synchronous changes in population dynamics. To test this hypothesis, we used *StairwayPlot2* to reconstruct the dynamics of population effective size change of the two most divergent clusters (YEN and EUR). Note that we did not transform the raw output from *StairwayPlot2* from generations to actual time, because of the uncertainties about generation time and mutation rate. Our results suggest high synchronicity patterns for both YEN and EUR populations; all populations show a similar trend for *Ne* change over generations (Fig. 4).

**Figure 4.**
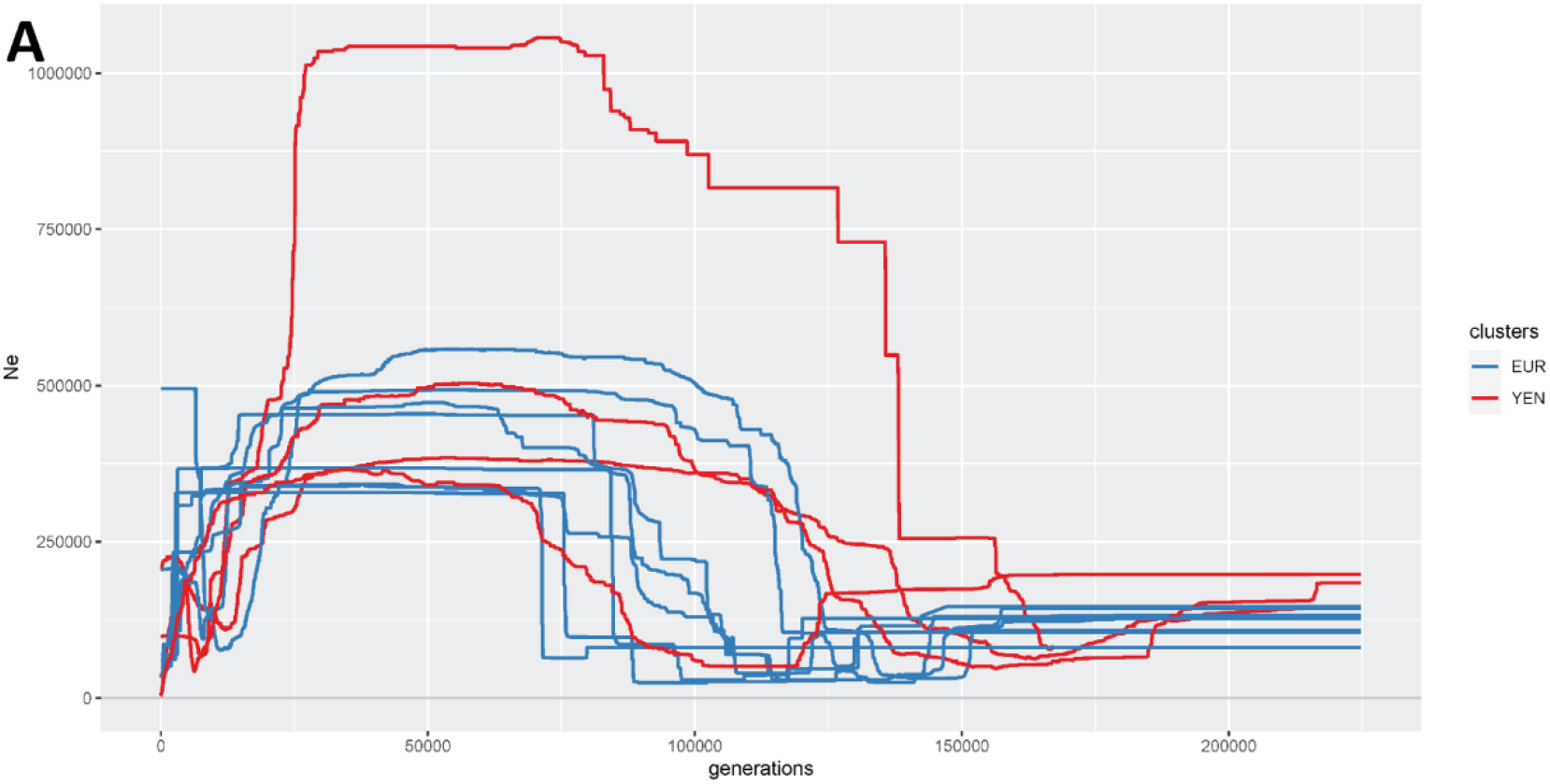
Change in Effective population size. Synchronicity of effective populations size (*Ne*) change through generations between populations from EUR (*P. abies*), in blue, and populations from YEN (*P. obovata*), in red, clusters.

## Discussion

In the present study, we used nuclear DNA and chloroplast DNA polymorphisms to investigate the joint demographic history of two dominant spruce species in Eurasian boreal forests, *P. abies* and *P. obovata.* Our results highlight three main features of this demographic history. Firstly, the majority of pivotal events occurred before the Last Glacial Maximum (LGM), indicating that the current population structure is not solely a result of post-LGM influences. Secondly, bidirectional gene flow/admixture played an extremely important role in shaping the geographical distribution of the current diversity. Thirdly, despite substantial gene flow occurring within the common habitat of these two species, the deep-rooted genetic structure persisted across generations. This remarkable resilience highlights the enduring impact of historical events on the genetic makeup of these species. Below we discuss the implications of these features on the inferences that can be drawn on the demographic history of the two species.

### ***Picea obovata* and *P. abies* population genetic structure differ**

We first used *UMAP* (PCA-based) and *ADMIXTURE* for genetic clustering. The PCA based approach is recommended for clustering, especially when genetic variation is distributed continuously across space, as it is the case for Norway and Siberian spruce (Novembre and Stephens 2008). In *P. abies*, our findings corroborate the primary population structure identified in earlier investigations (J. Chen et al. 2019; Li et al. 2022; Milesi et al. 2023). In particular, we confirmed that the genetic contribution of *P. abies* expands as far East as the Southern Ural Mountains (Tsuda et al. 2016, Fig. 2). Much less was known about *P. obovata* genetic structure and our study revealed the existence of at least three main genetic clusters: from east to west, a first cluster comprising populations sampled along the Yenisei River, a second one grouping populations along the Ob River, and finally a third one west of the Ural Mountains (Fig. 1 and S2&3). Strikingly, within each of those three clusters, each of which extends over long latitudinal gradients (up to 10 degrees between the southernmost and northernmost populations) there was almost no genetic differentiation. The distinct longitudinal structure detected in *P. obovata* may be attributed, at least in part, to the absence of samples from the expansive area between the Yenisei River and the Ob River. With a more comprehensive sampling in these regions, we might have observed a more gradual shift in allele frequencies, aligning with the broader patterns of Isolation-by-Distance (IBD) and our nucleotide diversity estimates. In any case, the longitudinal structure observed in *P. obovata*, is in strong contrast to *P. abies* where the main structure runs along latitude and mainly separate Southern clusters (ITA, EUR) from more northern ones (Fenno 1 and 2). This likely reflects the difference in glacial history between Western Europe and Siberia, where unlike Scandinavia, the latter was never fully glaciated (Velichko et al. 2011).

### ITA, PAH and Indigo are likely “relic” genetic clusters/populations

While the PCA and Bayesian inference-based clustering approach provided high resolution in the definition of today’s genetic clusters, *ADMIXTURE* analysis sheds light on their origin. These analyses are complementary and were largely consistent with each other, larger K in *ADMIXTURE* leading to the definition of the clusters identified with *UMAP* with few discrepancies, mainly ITA and PAH clusters, which on the other hand showed more consistency with Bayesian inference.

Out of the nine genetic clusters identified, ITA and PAH clusters included only few individuals from two close-by populations from Italy (ITAH and ITAL) and one high altitude population from the Southern Urals (PAH), respectively. ITAH and ITAL were collected from the southern limit of the natural range of Norway spruce and are geographically isolated from the other populations of the species, as confirmed by *fEEMS* analysis (Fig. 1 and Fig. S1). The low nucleotide diversity of these populations further supports a higher level of genetic drift than that in the other populations of *P. abies*. Similarly, their high Tajima’s D estimates compared with that of the other populations tend to show that their demography histories also differ from that of the surrounding populations. Within *P. obovata*, individuals from PAH, a population located at high altitude, clustered separately from all nearby populations (PCA-UMAP); including PAL, a population of low altitude at the same location. In contrast, PAH is closer than expected given geographic distance to Indigo population which is located at the very northern range of *P. obovata* (Fig. 2B and Fig. S3). Both PAH and Indigo share phenotypic traits characteristic of adaption to harsh habitats with extreme temperatures and strong winds; typically, small trees with a “bushy” architecture (V. Semerikov and M. Lascoux, personal observation), as also encountered at high altitude in Norwegian and Swedish mountain ranges (for a picture, see Nota et al. 2022). The observed pattern could thus be a result of convergent local adaptation to extreme environment. A more likely explanation, though not exclusive, is that both populations survived through glacial periods as small and isolated refugia and received limited influence from any other populations since then. The closer-than-expected relationship could thus also reflect a shared ancestry and isolation. Sediment coring analysis confirmed the presence of spruce pollen during LGM in Indigo (Väliranta et al. 2011). It is well known that high mountains served as refugia for Norway spruce populations during glacial periods (Nota et al. 2022; Parducci et al. 2012; Tollefsrud et al. 2008, 2015) and the same probably applies to the PAH population. In Fennoscandia, however, these high-altitude relics did not have a significant contribution to the re-colonization after the LGM (Nota et al. 2022). Interestingly, the allelic difference between PAH and the nearby PAL also indicates a limited contribution of these relict populations to post-glacial recolonization, which is not surprising considering the fact that trees of PAH are small and seem to primarily reproduce asexually (V. Semerikov and M. Lascoux, personal observation). Similar to ITA, PAH and Indigo are characterized by a low nucleotide diversity when compared with surrounding populations and are “pure” genetic clusters. It is thus likely that PAH and Indigo in *P. obovata*, as well as ITA in *P. abies* are somehow the relics of past genetic distribution.

### **Two ancient cryptic refugia bridge *P. abies* and *P. obovata* genetic background**

Recent literature suggested a pattern of introgression of *P. abies* populations by *P. obovata* genetic background, in particular in the Northern range of *P. abies* (J. Chen et al. 2019; Li et al. 2022; Tollefsrud et al. 2015; Tsuda et al. 2016). The analyses of mitochondrial DNA and of 10 SSR markers by Tsuda et al. (2016) also suggested that cryptic *P. abies* populations might have acted as stepping stones for Siberian spruce westwards move. The presence of those “stepping stones” would help explain the asymmetric patterns of introgression observed for mtDNA and nuclear DNA (Tsuda et al. 2016) and would also be consistent with Currat et al. (2008) introgression model. In line with these results, the admixture analysis suggested that K= 3 or K=4 ancestry components were at the origin of today’s population structure (Fig. 2B). It implies that at least one (K = 3), or more probably two (K = 4), ancestry components contributed to the genetic variation we observe today. Additionally, we detected four main chloroplast haplotypes. Their phylogenetic relationships and spatial distribution suggest that these haplotypes arose from the same ancestral populations as the ones identified using nuclear polymorphisms. This further supports our hypothesis of the existence of two cryptic refugia, one with a more *P. abies* genetic background (AC_FEN and CP group2, Fig. 2) and one with a more *P. obovata* background (AC_UR and CP group3, Fig. 2).

However, previous studies identified only three main mitochondrial groups across the distribution of the two species using the gene *nad1* as mitochondrial marker (Tollefsrud et al. 2008, 2015; Tsuda et al. 2016) where we could have expected four mitochondrial groups if the various ancestry component originated from completely isolated lineages. Among them, two haplotype groups were restricted to *P. abies* (Central Europe, Alpine and Carpathian Mountains) or *P. obovata* (East of the Ural Mountains). The last one covered the whole hybrid zone and expanded from the Ural Mountains in the East, to the Baltic countries and the whole Fennoscandia in its Western range (see Fig. 1 in Tsuda et al. 2016). Despite their current distribution, these haplotypes are more closely related to those only found in *P. obovata* than those only found in *P. abies*. Although one can’t rule out the possibility of a lack of resolution of the mitochondrial maker used (< 1000 bp), a more likely explanation lies in the different mode of inheritance and dispersion of mitochondrial DNA (maternally inherited, only dispersed through the seeds), of chloroplast DNA (paternally / seed and pollen) and of nuclear DNA.

Combining nuclear and chloroplast data from our study with the distribution of mitochondrial DNA from Tsuda et al. 2016, we can infer that past populations in the hybrid zone were likely founded by westward colonization from the *P. obovata* domain, the mitochondrial haplotype being retained into the two refugia in the hybrid zone. Individuals from one refugium then received a massive gene flow from *P. abies* through pollen dispersal and the local chloroplast genome was replaced by the one from Norway spruce (chloroplast capture, Schulte et al. 2021; Tsitrone, Kirkpatrick, and Levin 2003; Yang et al. 2021) while the other refugium was more influenced by *P. obovata*. This hypothesis would also explain the higher divergence between CP group2 and CP group3 than expected when considering the low *F_ST_* between the corresponding ancestral components estimated with *ADMIXTURE* (Tab. 3). Both, a tree-based approach (*TreeMix*) and coalescent simulations (*FastSimCoal2*) supported recurrent admixture events between the main genetic entities that gave rise to the main genetic clusters that survived glacial periods. Even if a precise timing of these admixture events is out of reach given the uncertainties about mutation rate and generation time in spruce species, it is worth noting that they all date back before the LGM. Our best estimates showed that the main admixture events between *P. abies* and *P. obovata* genetic background occurred during the Pleistocene in agreement with what was already described (J. Chen et al. 2019; Tsuda et al. 2016) and likely during inter-glacial recolonization. Note that a direct comparison of the estimates between these studies would be meaningless as the demographic models and the gene pool considered are different in the different studies.

**Table 3.**
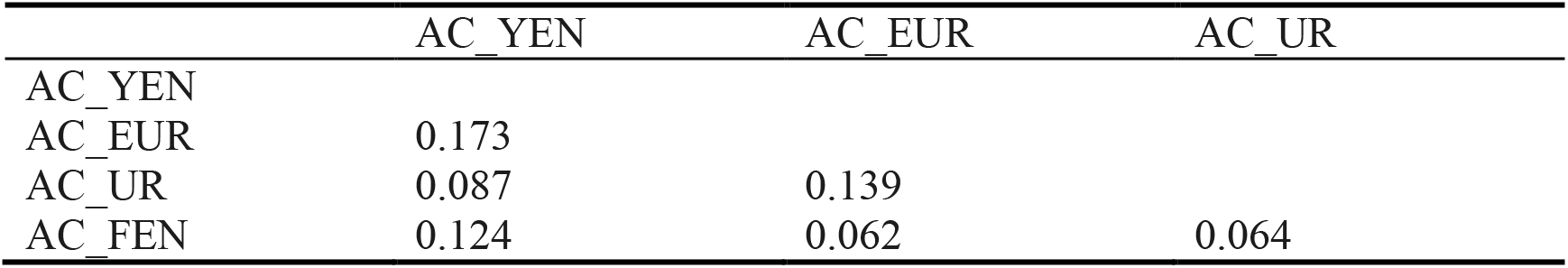
Genetic differentiation among the putative ancestry populations.

The location of these refugia remains unclear but our analyses points toward Western Russia-Eastern Fennoscandia and the Southern Ural Mountains. Tollefsrud et al. (2008) showed that early Holocene pollens were enriched in the Alps and Carpathian Mountains, which are known refugia for Norway spruce, but also in a large area in Western Russia. The latter, could thus be the location of one refugium during glacial periods as also suggested by Giesecke and Bennett (2004) and Latałowa and van der Knaap (2006). Locating the other refugium could prove to be more challenging as the classical concept of refugia may not apply due to the lack of complete glaciation of the region. According to paleobiological records Siberia was a dry desert interspersed with pockets of forested areas (Semerikov et al. 2013 and reference there in). Still, Karunarathne et al. (2023) suggested that the southern part of the Siberian Plain was a suitable habitat for Siberian spruce during glacial periods and the presence of spruce during the LGM in the Baikal region is attested by fossil pollen (Kobe et al. 2022). Using niche modelling and backward climate projection, Karunarathne et al. (2023) also suggested that the distribution ranges of *P. abies* and *P. obovata* overlapped during interglacial periods in their Northern range (∼145 kya). This could for instance explain both the admixture of the two species at the origin of the Fenno1 cluster (∼180 kya, Fig. 3B) and the fact that *P. obovata* contribution expands further West in the Northern range of *P. abies*.

### Local adaptation probably maintained genetic clusters under extensive gene flow

Today’s genomic variation shows a continuous spatial distribution between *P. obovata* and *P. abies*. The low differentiation between populations geographically distant and belonging to different genetic clusters having diverged long ago highlights the extensive gene flow and or migration occurring between the various genetic entities (Fig 2B, S3). The gene flow is sustained by long range pollen dispersal, as evidenced by the distribution of the chloroplast haplotypes when compared to the distribution of nuclear DNA variation. Despite such a high gene flow, well delineated genetic clusters persist at the nuclear level, probably because of the low dispersion of seeds and of the recurrent sheltering by well identified refugia during glacial periods. While some well identified barriers to gene flow explain the distribution of current genetic diversity (e.g. mountain ranges, the Baltic Sea), no obvious topographic barrier can explain the main barrier running within the hybrid zone. This barrier could thus be the result of the interplay of the ancient demographic events described above with ecological factors and local adaptation. A whole body of literature actually discriminates *P. abies* from *P. obovata* and their hybrids based on the shape of the cone scale (e.g. Pravdin 1975; Popov 2003, 2010) and their distribution match the distribution of the main genetic clusters within the hybrid zone we described. Two types of hybrids can be further discriminated using morphological traits, hybrids with properties of *P. abies* or with properties of *P. obovata* (e.g. E. Nakvasina et al. 2019; Orlova et al. 2020) mirroring the distribution of the ancestry components within the hybrid zone. Recently, Nakvasina et al. (2019) showed that the various hybrid forms display different phenotypic plasticity and that both hybrid forms had higher survival rates than the non-hybrids. The provenance test was conducted in the Arkhangelsk Region, the highest survival rate being achieved by the local genotype, namely hybrid with “*P. obovata*” properties. Similarly, Li et al. (2022) have shown that the contact zone between Fenno1 and Fenno2 clusters in Sweden matched the transition between main climatic zones and they evidenced the role of local adaptation in the maintenance of the contact zone despite extensive gene flow. These studies thus further support the role of natural selection acting in maintaining the hybrid zone between locally adapted ecotypes, what would be addressed in another manuscript.

Far from limiting the gene flow between the two species, it seems that recurring hybridization events between *P. abies* and *P. obovata* gave rise to different genetic entities that survived ice-age. Their recent contact after LGM is probably at the origin of locally adapted ecotypes structuring the large hybrid zone between the two species. In line with this hypothesis, Karunarathne et al. (2023) showed that the hybridization between the two species enlarged both species ecological niches, hybrids occupying a specific ecological niche.

## Conclusion

This study showed that the current distribution of genetic diversity of *P. abies* and *P. obovata* was mainly shaped by a complex demographic history involving repeated hybridization events between Norway and Siberian spruce during the Pleistocene. Two cryptic refugia in the large hybrid zone probably survived the glacial periods and played a critical role in shaping the current distribution of the two species of spruce. Instead of only an introgression of *P. abies* populations by *P. obovata*, our study suggests a two-pronged introgression pattern, with *P. obovata* expanding into *P. abies* Northern range and *P. abies* introgressing *P. obovata* populations in their southern range. Recent genomic studies revealed that hybridization and admixture are common in many forest trees species and our study highlights the importance of considering the whole species complex instead of separate entities to shed light on their complex demographic histories.

## Supporting information

Supp1: model comparison

Supp2: individual information

Supp3: population information

Supp4: results for f3 test

Supp5: goodness of fit evaluation

## Acknowledgements

We would like to thank Swedish National Infrastructure for Computing (SNIC) for resource allocation for high-power computing and data storage under the project numbers SNIC 2022/6-313 and SNIC 2022/5-543. We also would like to thank Vladimir Semerikov for his help with sampling most of the *P. obovata* material and in providing us with additional plant materials. The Lundman’s Foundation for Botanical studies grant from the Swedish Phytogeographic Society awarded to Qiujie Zhou was immensely helpful for covering the cost for high-throughput sequencing.

