## Supplementary figures for "Recurrent hybridization and gene flow shaped Norway and Siberian spruce evolutionary history over multiple glacial cycles"

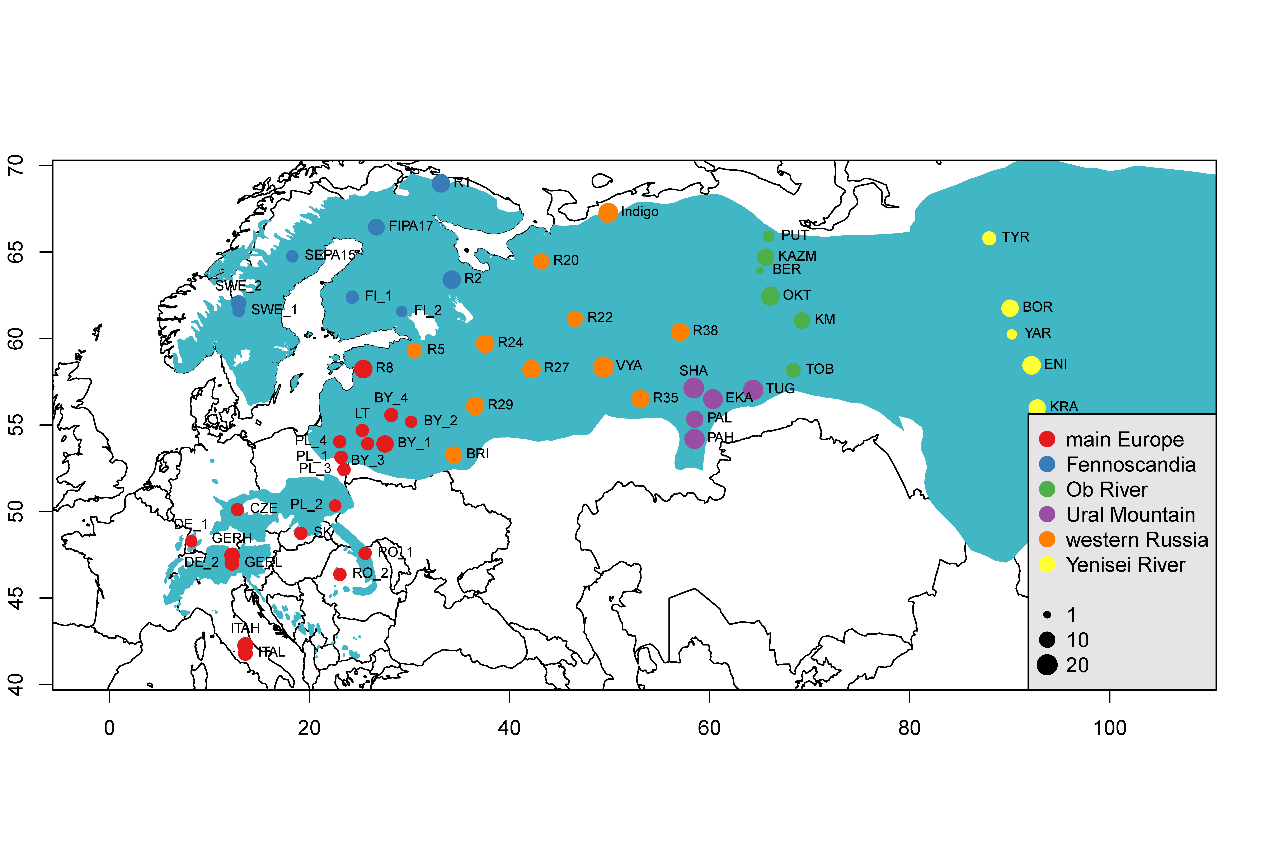

**Figure S1**. Populations from *Picea abies*, *P. obovata* and putative hybrids/ introgressed individual sampled in this study. Blue shadow shows the joint distribution of *P. abies – P. obovata* complex. The red dotted lines indicate roughly the proposed longitudinal limit of the hybrid zone between Norway spruce and Siberian spruce.

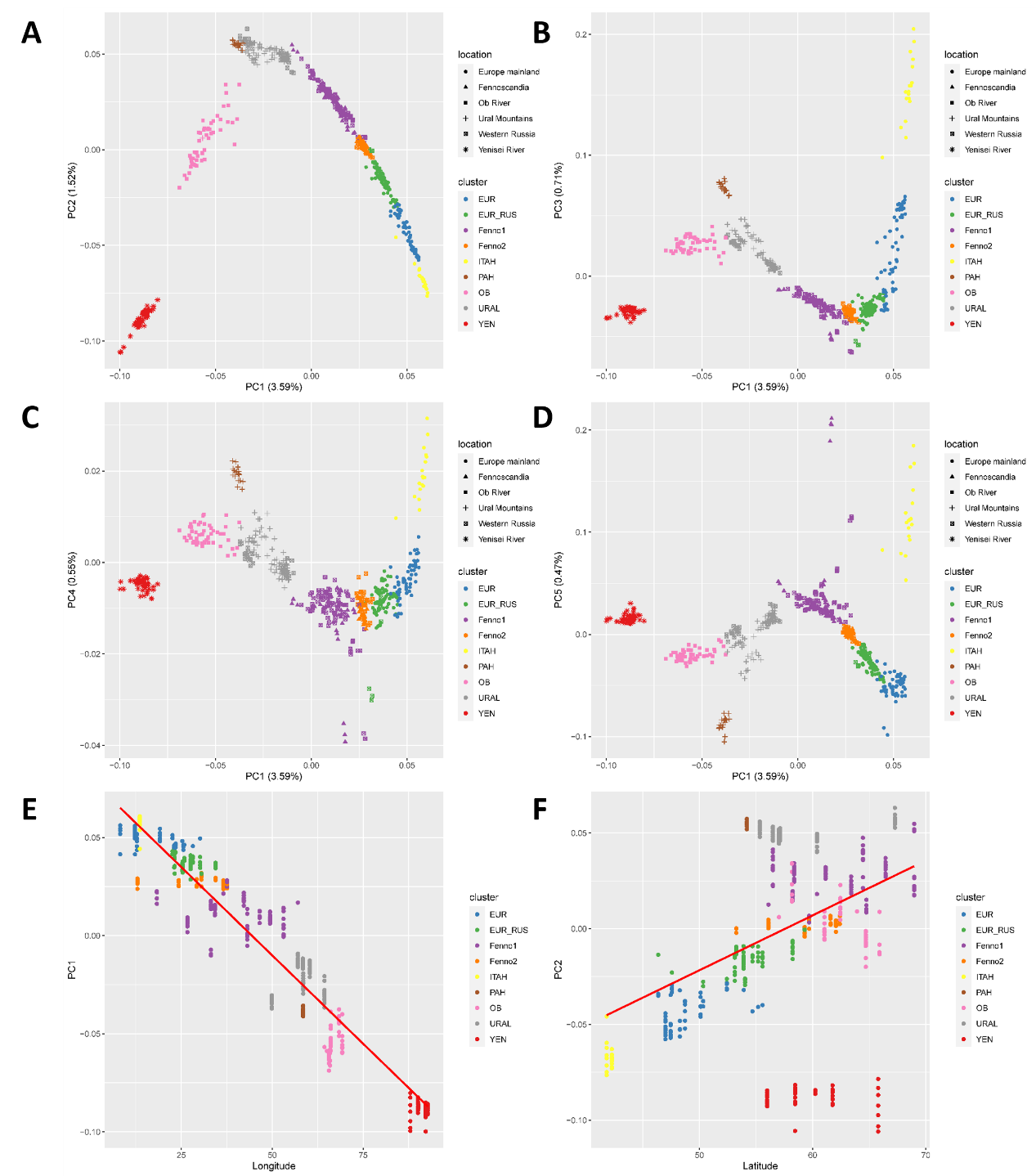

**Figure S2**. Population structure revealed by principal component analysis (PCA). A, B, C, D, results of PCA shown as plotting PC2, PC3, PC4 and PC5 against PC1, respectively; E. regression PC1 over longitude of each individual; F, regression PC2 against latitude of each individual. Individuals were colored with its assigned genetic clusters.

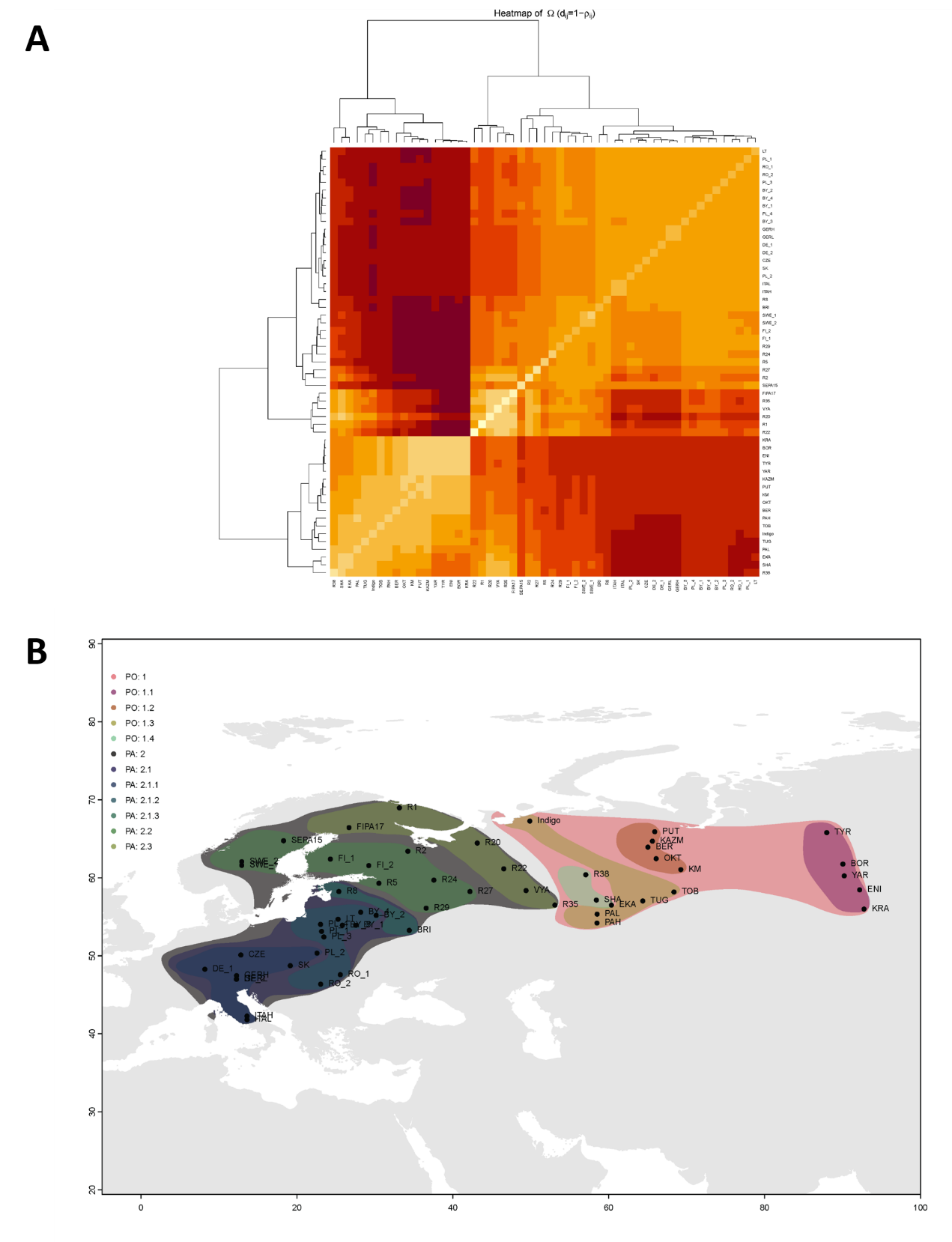

**Figure S3**. Genetic clustering at population level inferred with Bayesian inference. A, Heatmap showing the covariance of Bayesian factor among populations. Clustering of populations of *P. abies* and *P. obovata* based on Bayesian inferences, the genetic distance matrix assumes the allele frequencies of populations diverge from a hypothetical allele frequency due to drift and hence some populations can covary due to shared demographic history and gene flow (Coop et al. 2010). B, geographic projection of the inferred genetic clusters.

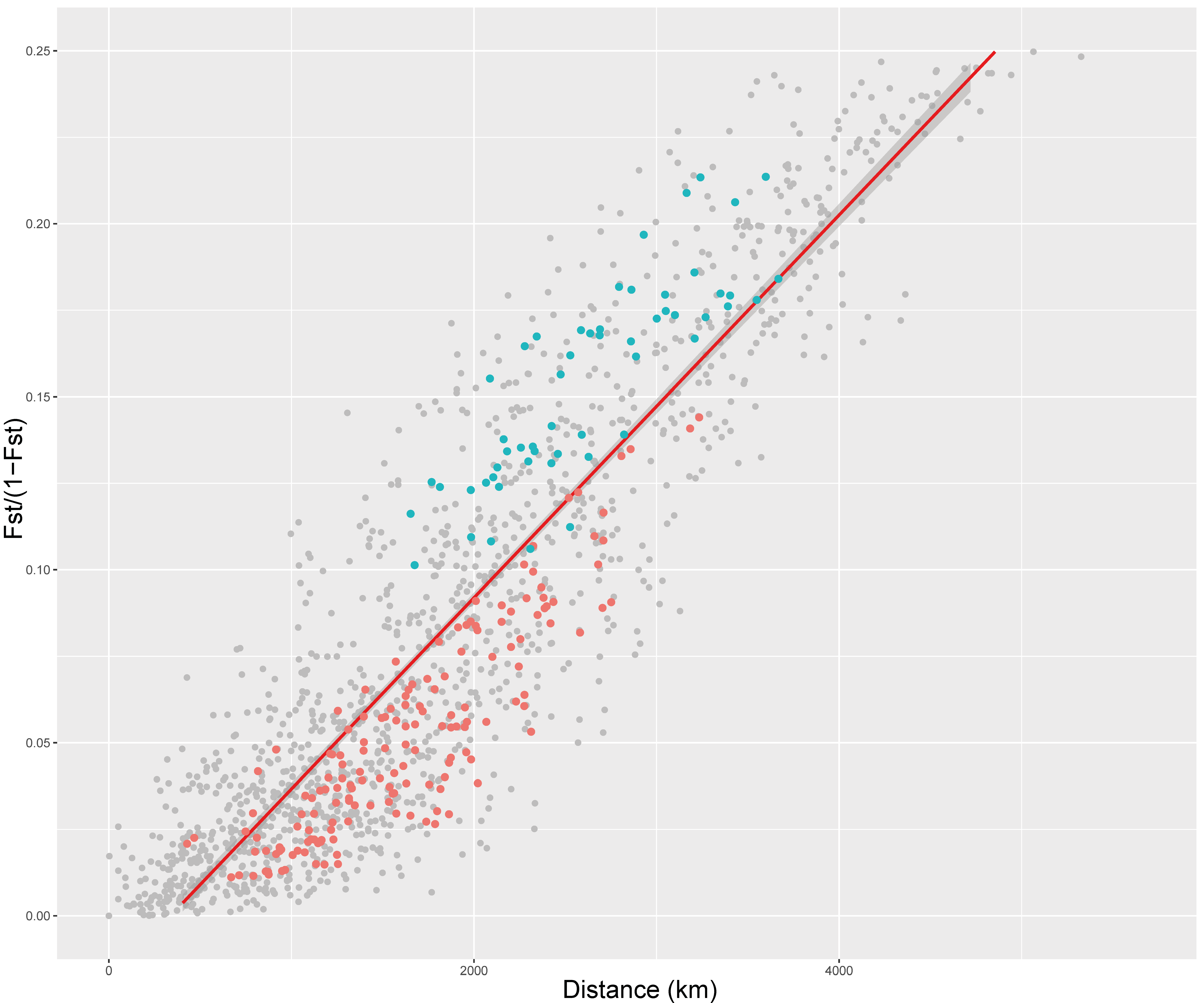

**Figure S4**. Pattern of isolation by distance shown as regression *Fst*/(1-*Fst*) over geodesic distance for each population pair. Blue points represent populations pairs consisting of one population from Western Russia and the other from Yenisei River while red points show populations pairs consisting of one population from main Europe and the other from Fennoscandia. See Fig S1 for population assignation to different geographic regions.

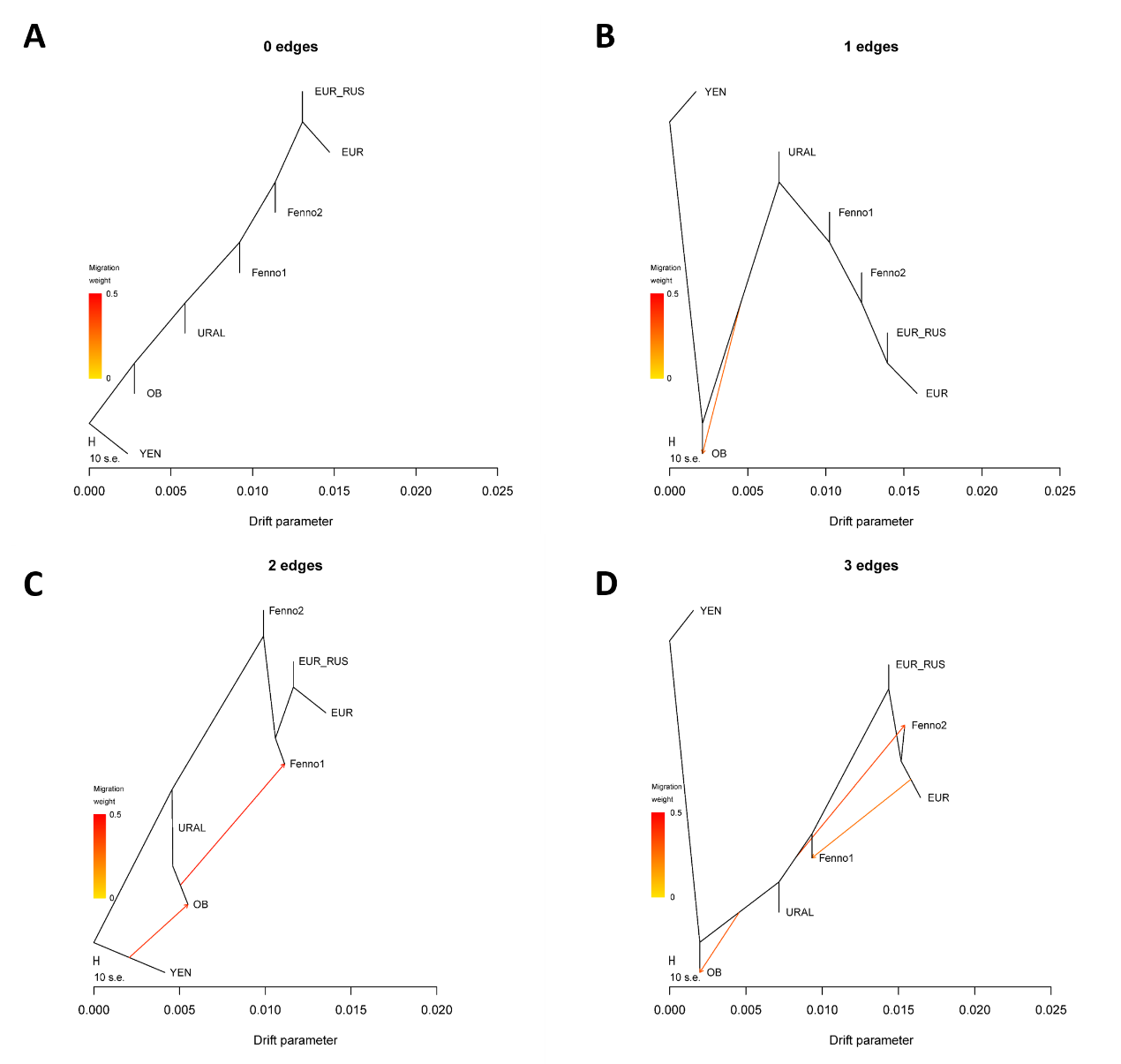

**Figure S5**. Admixture events inferred with *TreeMix*. A, B, C and D, evolutionary scenarios suggested by *TreeMix*, with 0—3 edges added to the inferred maximum likelihood phylogenetic tree, respectively.
