## Supplementary material for "Recurrent hybridization and gene flow shaped Norway and Siberian spruce evolutionary history over multiple glacial cycles": Supp5: goodness of fit evaluation

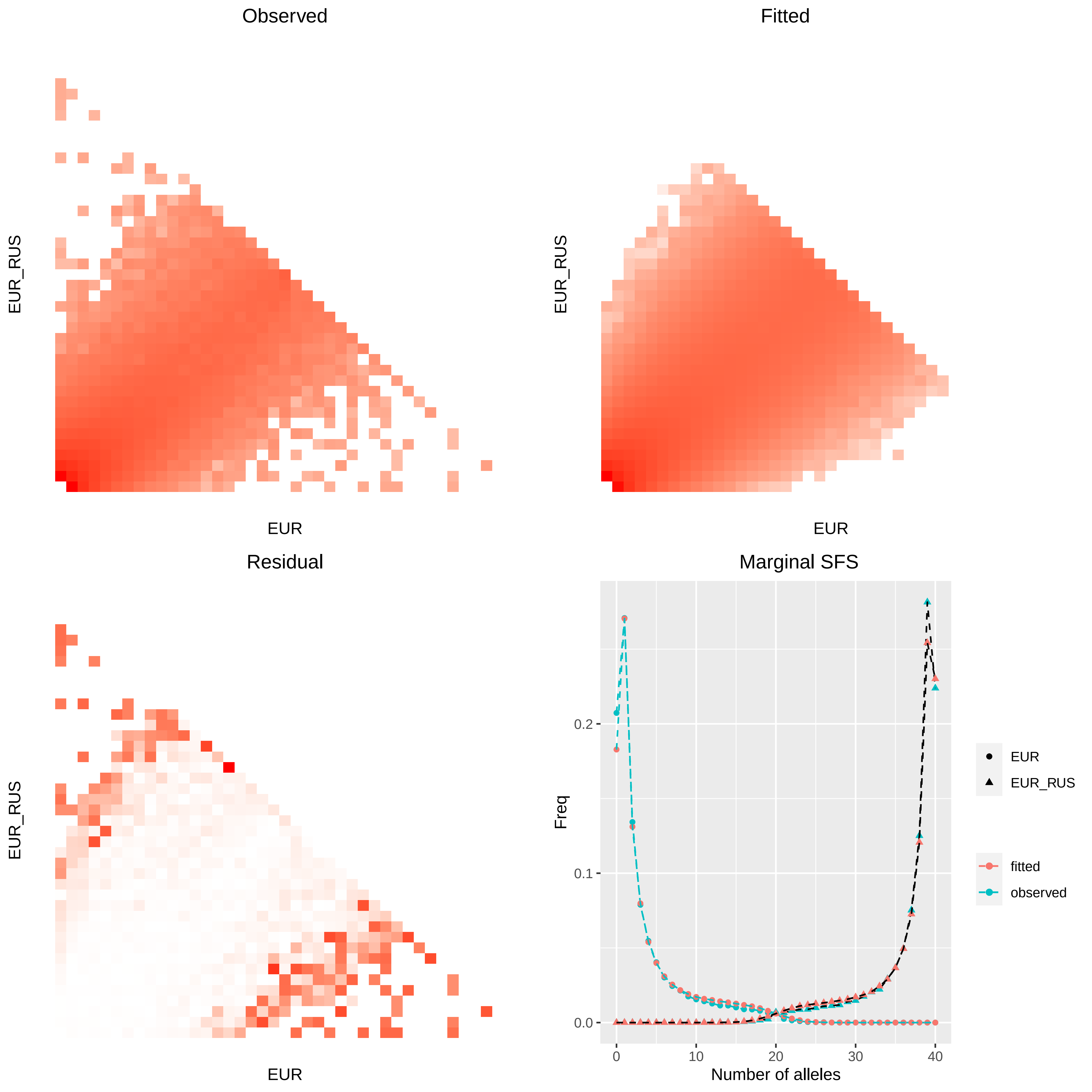
Observed
EUR_RUS
EUR
Fitted
EUR_RUS
EUR
Residual
EUR_RUS
EUR
Marginal SFS
Freq
0.0
0.1
0.2
0
10
20
30
40
Number of alleles
EUR
EUR_RUS
fitted
observed

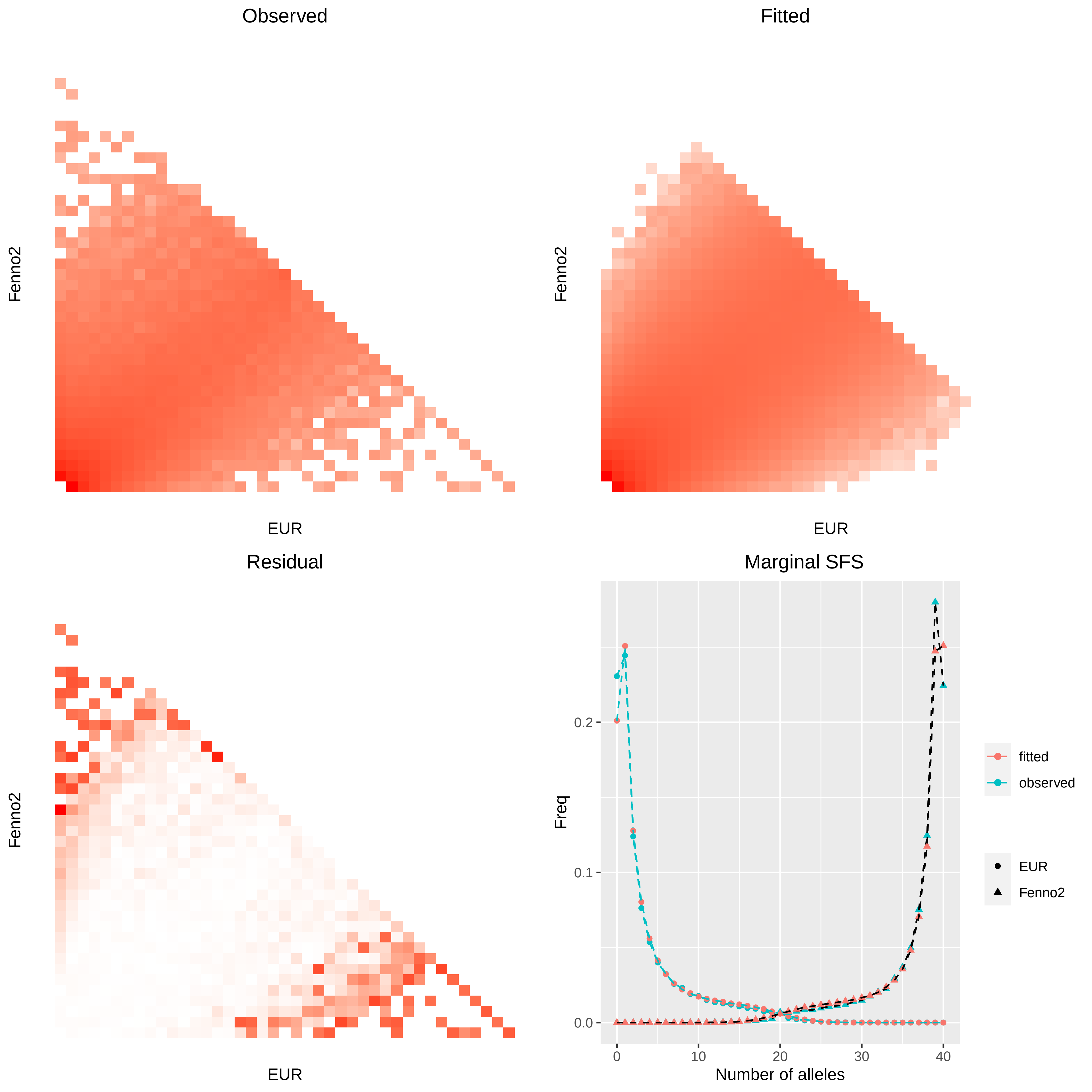

Observed
Fenno2
EUR
Fitted
Fenno2
EUR
Residual
Fenno2
EUR
Marginal SFS
Freq
0.0
0.1
0.2
0
10
20
30
40
Number of alleles
fitted
observed
EUR
Fenno2

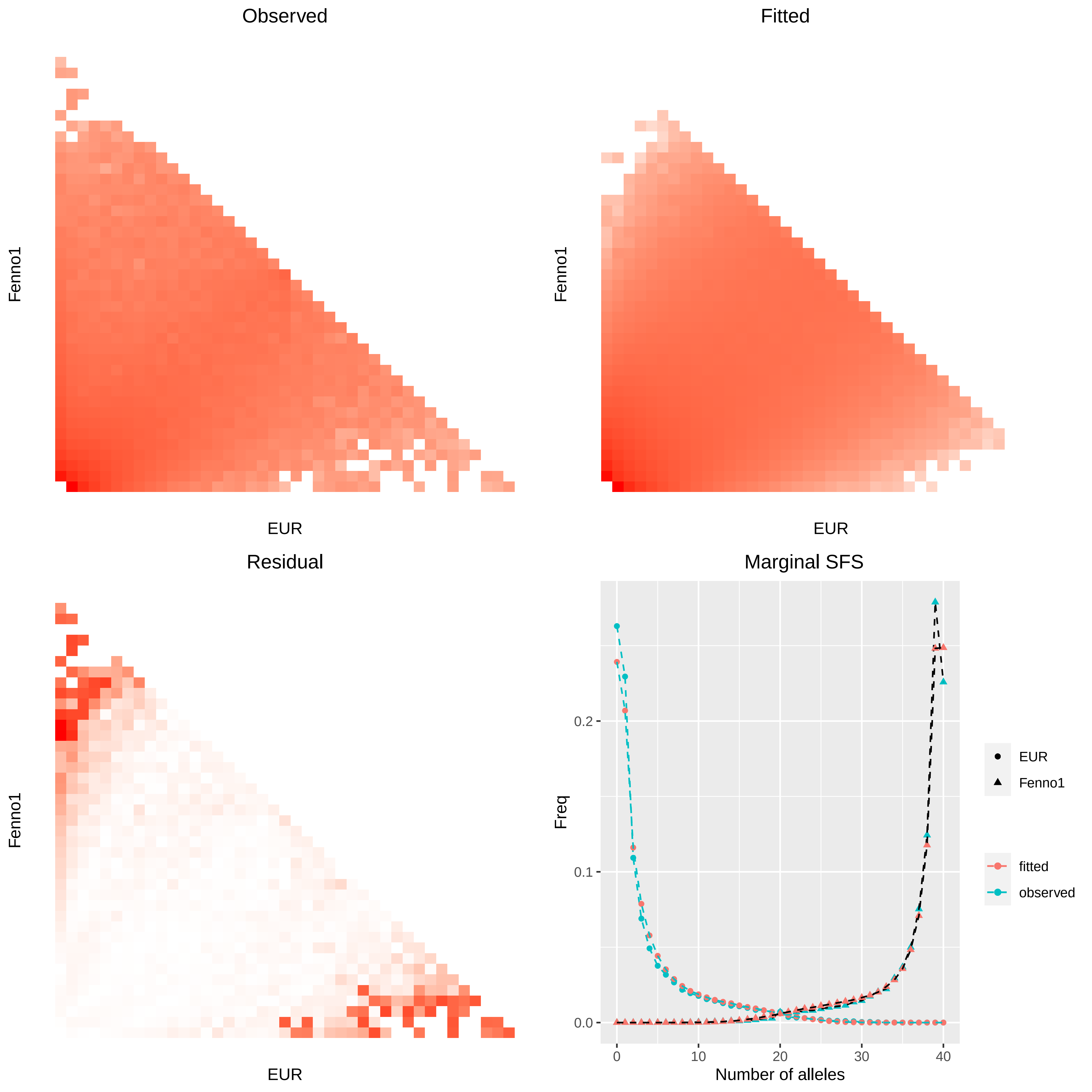

Observed
Fitted
Residual
Marginal SFS
Fenno1
EUR
Freq
Number of alleles
0.0
0.1
0.2
0
10
20
30
40
EUR
Fenno1
fitted
observed

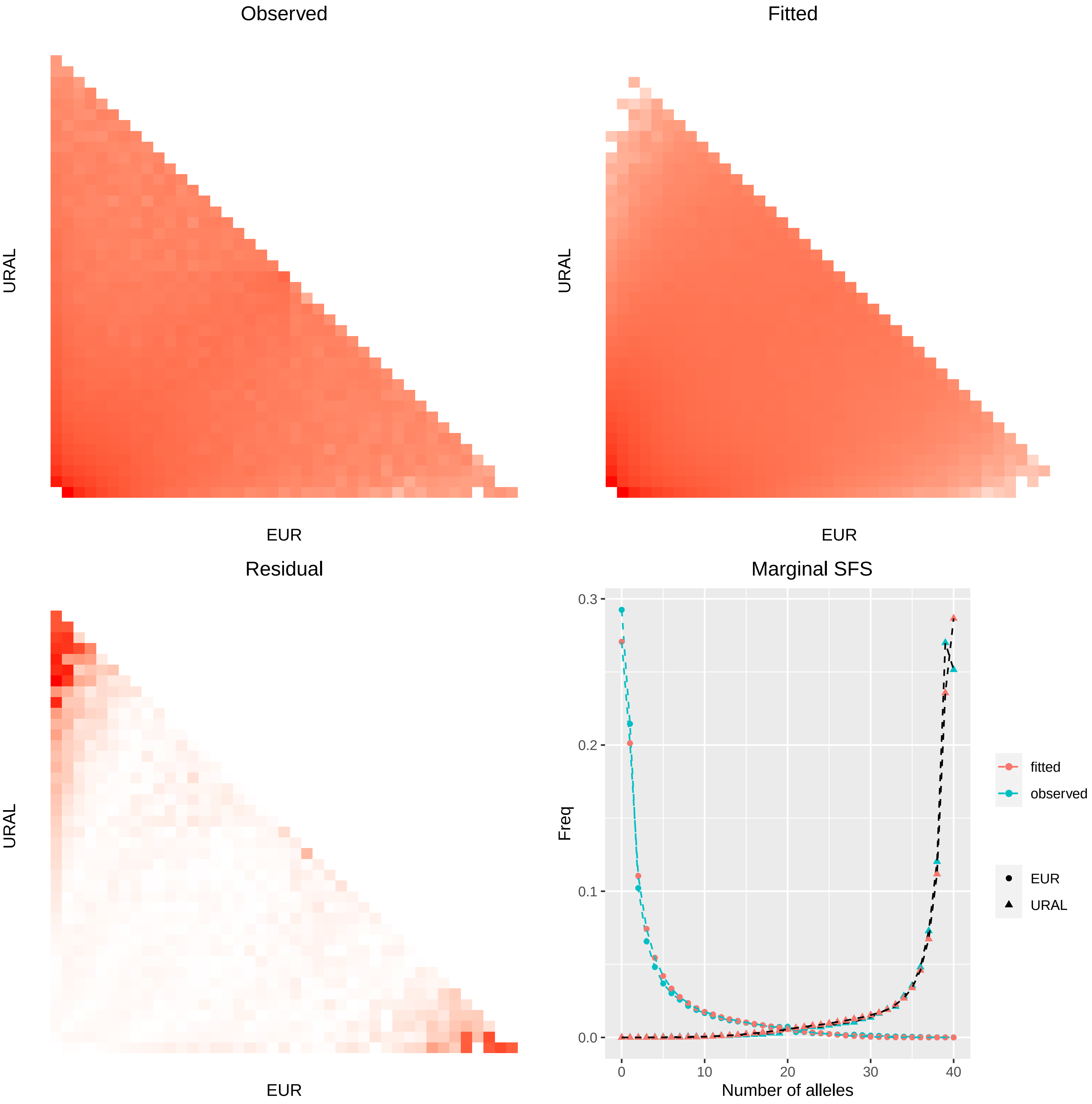

Observed
URAL
EUR
Fitted
URAL
EUR
Residual
URAL
EUR
Marginal SFS
Freq
0.3
0.2
0.1
0.0
0
10
20
30
40
Number of alleles
fitted
observed
EUR
URAL

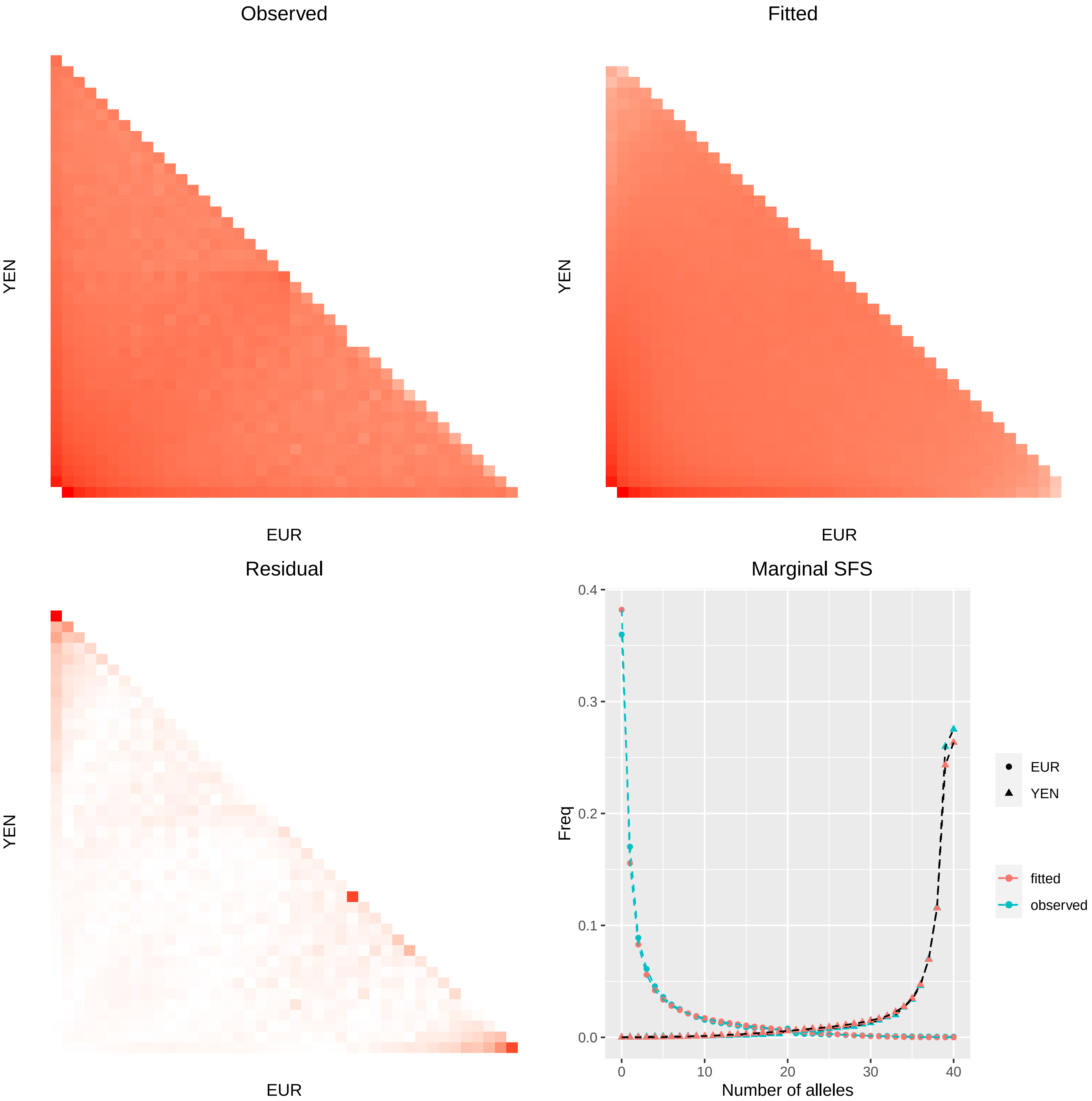
Observed
YEN
EUR
Fitted
YEN
EUR
Residual
YEN
EUR
Marginal SFS
Freq
0.4
0.3
0.2
0.1
0.0
0
10
20
30
40
Number of alleles
EUR
YEN
fitted
observed

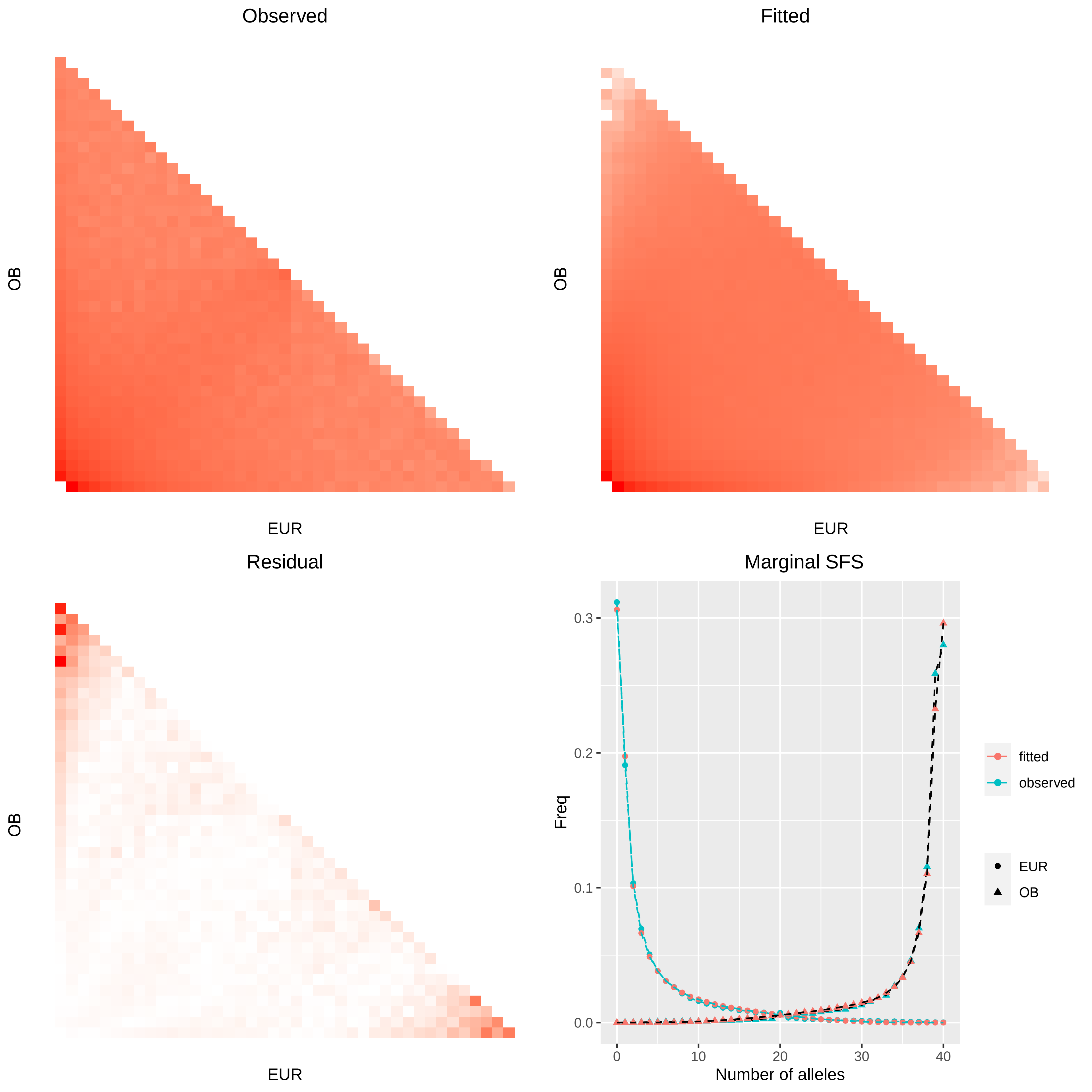

Observed
OB
EUR
Fitted
OB
EUR
Residual
OB
EUR
Marginal SFS
Freq
0.0
0.1
0.2
0.3
0
10
20
30
40
Number of alleles
fitted
observed
EUR
OB

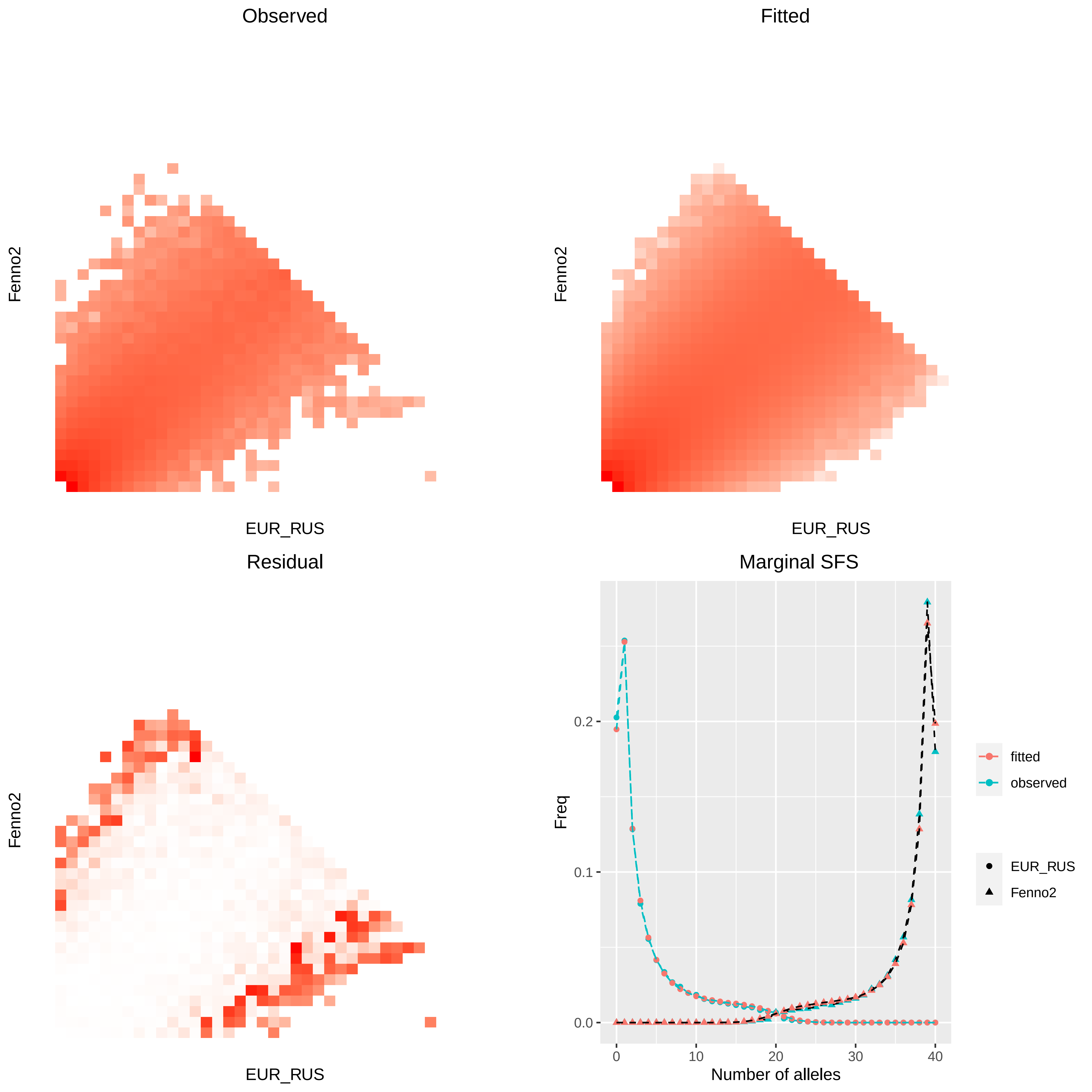

Observed
Fitted
Residual
Marginal SFS
Fenno2
EUR_RUS
Freq
Number of alleles
0.0
0.1
0.2
0
10
20
30
40
fitted
observed
Fenno2

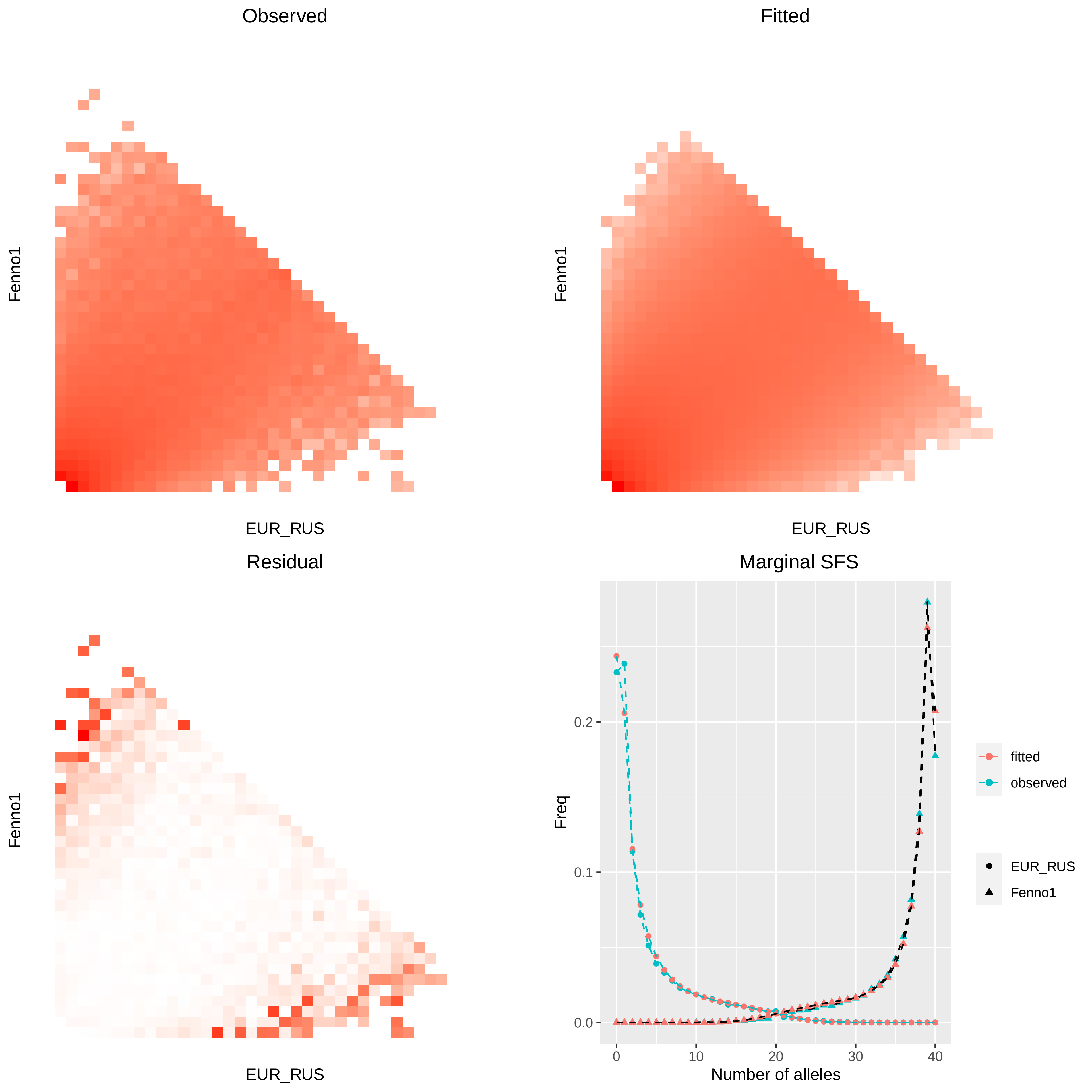

Observed
Fitted
Fenno1
EUR_RUS
Residual
Marginal SFS
Freq
0.0
0.1
0.2
0
10
20
30
40
Number of alleles
fitted
observed
EUR_RUS
Fenno1

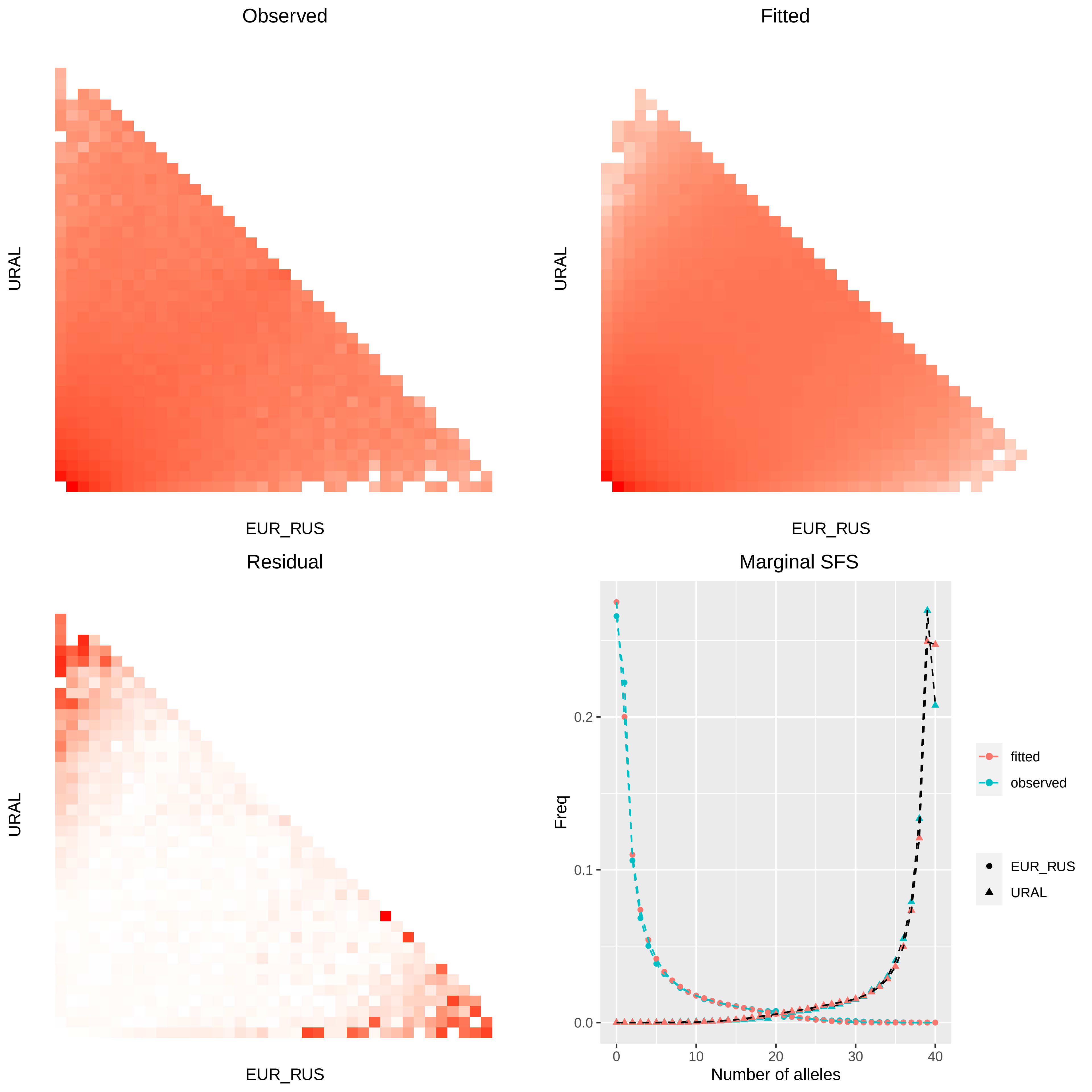

Observed
Fitted
URAL
EUR_RUS
Residual
Marginal SFS
Freq
0.0
0.1
0.2
0
10
20
30
40
Number of alleles
fitted
observed
EUR_RUS
URAL

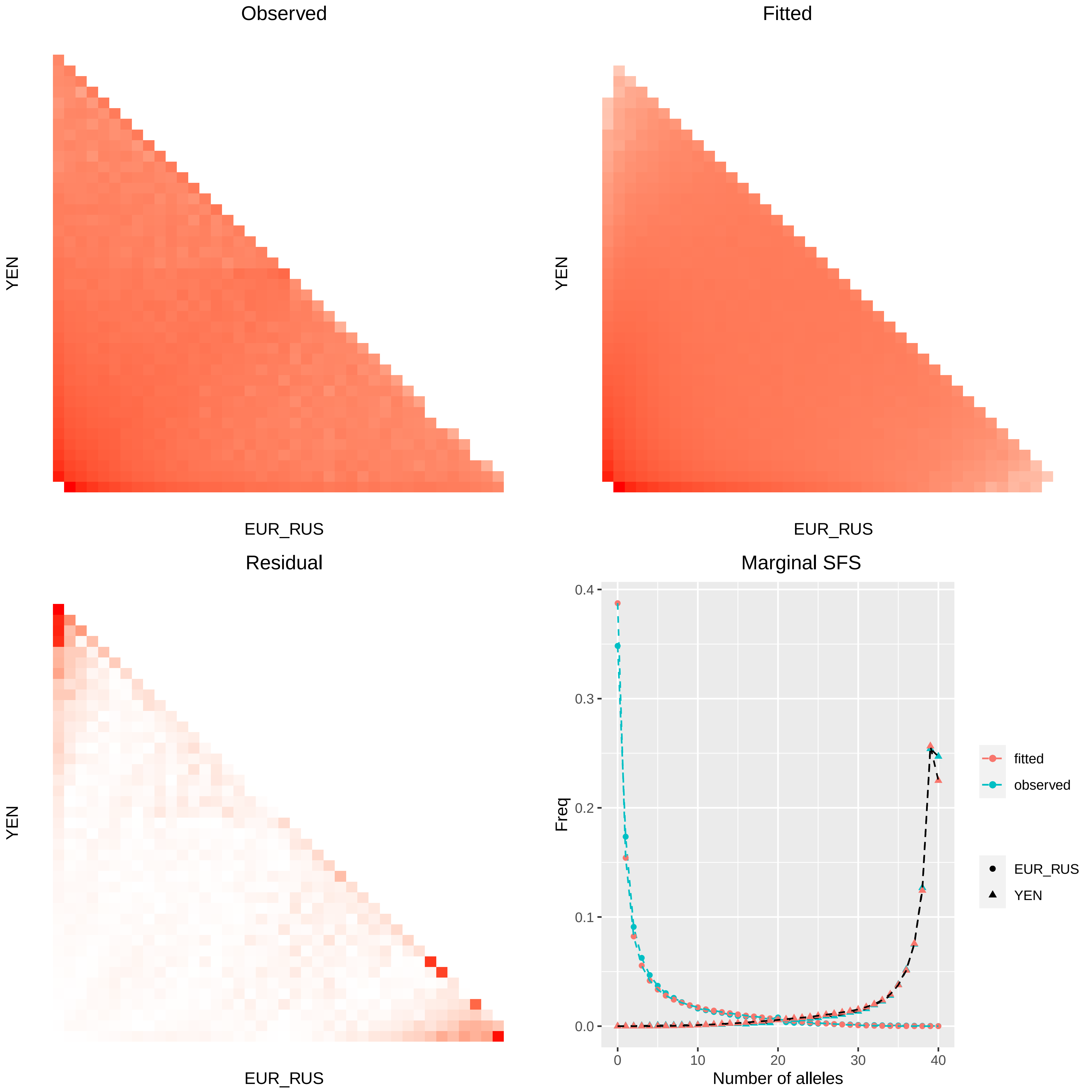

Observed
YEN
EUR_RUS
Fitted
YEN
EUR_RUS
Residual
YEN
EUR_RUS
Marginal SFS
Freq
0.0
0.1
0.2
0.3
0.4
0
10
20
30
40
Number of alleles
fitted
observed
EUR_RUS
YEN

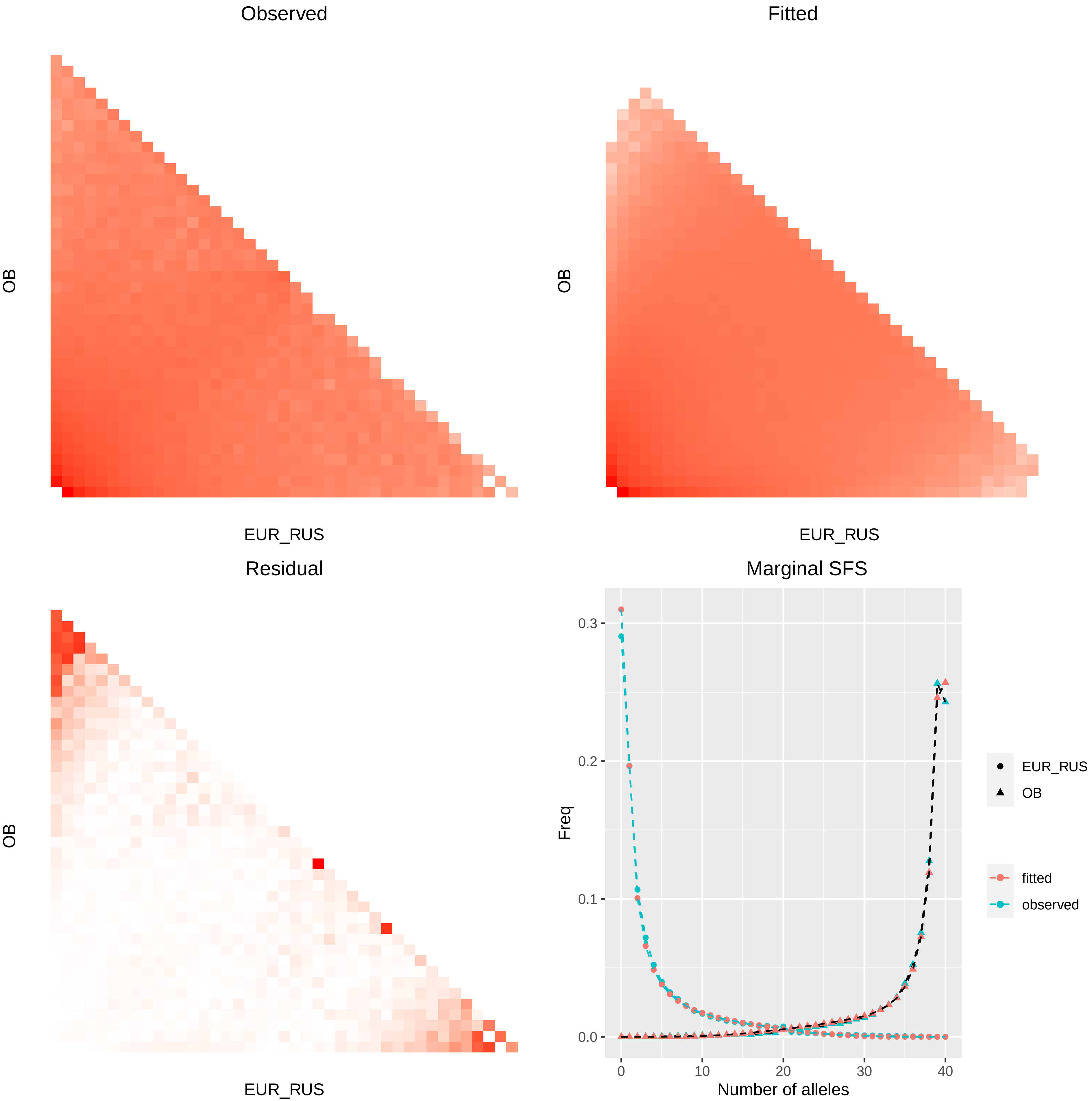
Observed
OB
EUR_RUS
Fitted
OB
EUR_RUS
Residual
OB
EUR_RUS
Marginal SFS
Freq
0.3
0.2
0.1
0.0
0
10
20
30
40
Number of alleles
EUR_RUS
OB
fitted
observed

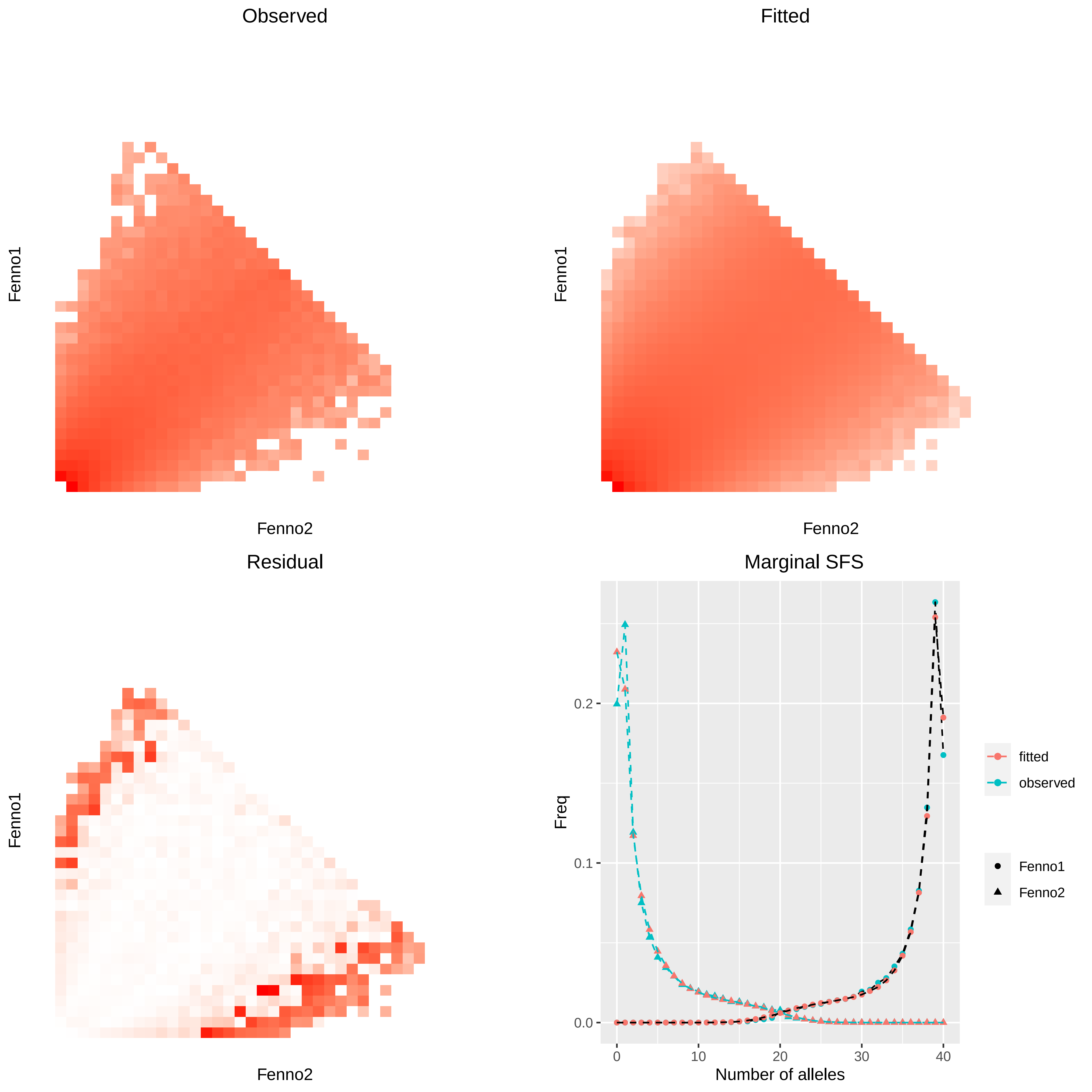

Observed
Fenno1
Fenno2
Fitted
Fenno1
Fenno2
Residual
Fenno1
Fenno2
Marginal SFS
Freq
0.0
0.1
0.2
0
10
20
30
40
Number of alleles
fitted
observed
Fenno1
Fenno2

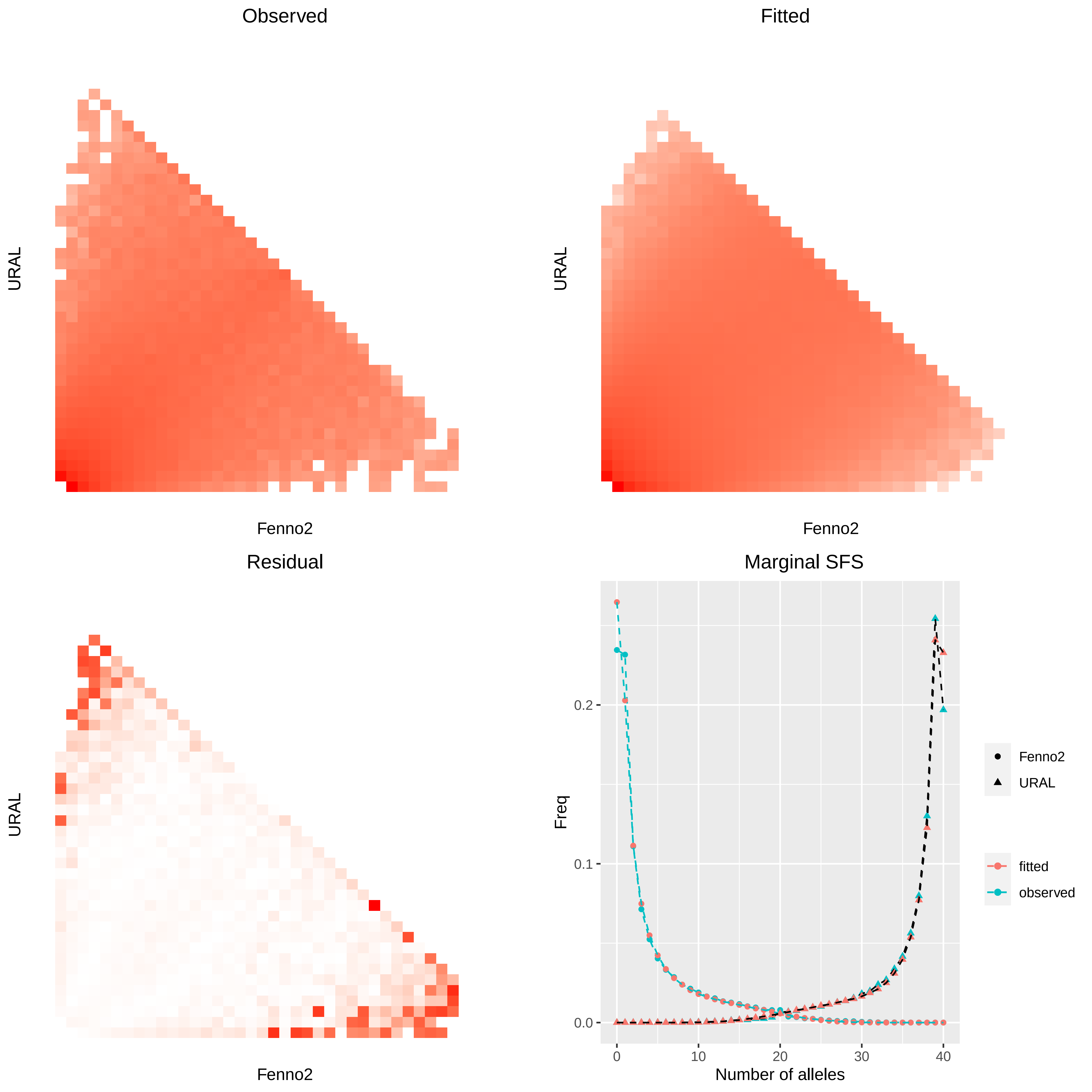
Observed
URAL
Fenno2
Fitted
URAL
Fenno2
Residual
URAL
Fenno2
Marginal SFS
Freq
0.0
0.1
0.2
0
10
20
30
40
Number of alleles
Fenno2
URAL
fitted
observed

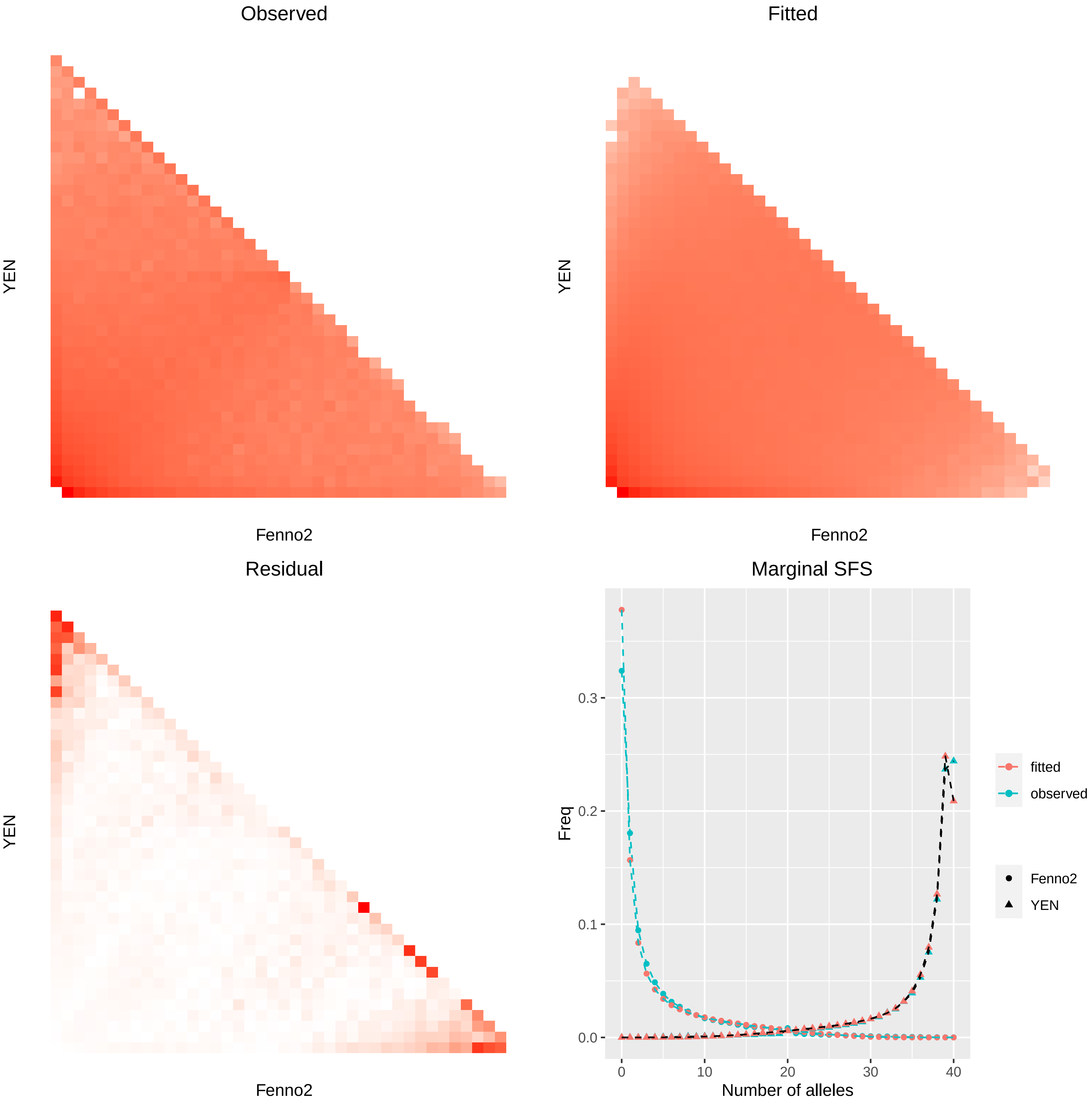
Observed
YEN
Fenno2
Fitted
YEN
Fenno2
Residual
YEN
Fenno2
Marginal SFS
Freq
0.0
0.1
0.2
0.3
0
10
20
30
40
Number of alleles
fitted
observed
Fenno2
YEN

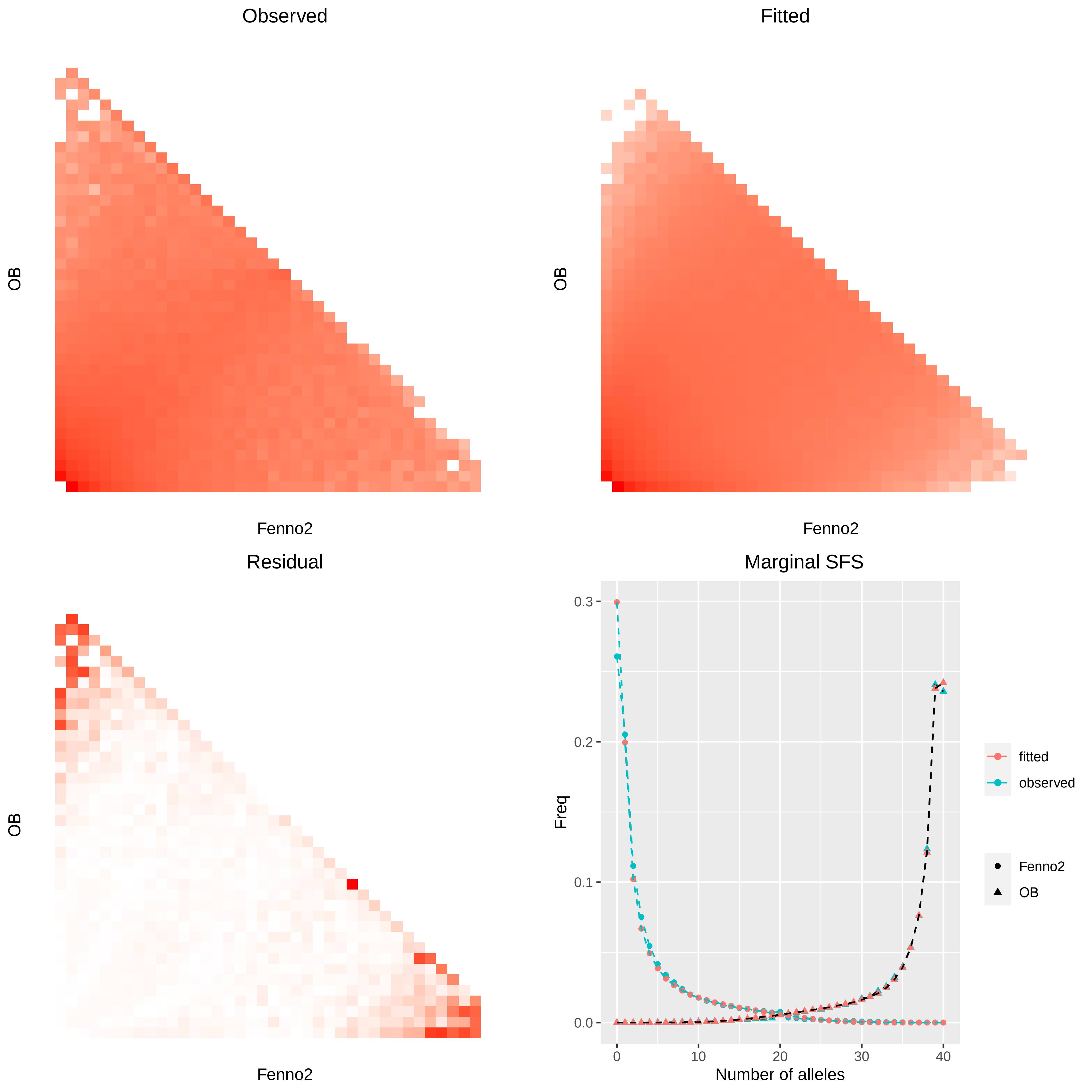
Observed
Fitted
Residual
Marginal SFS
OB
Fenno2
Freq
Number of alleles
0.0
0.1
0.2
0.3
0
10
20
30
40
fitted
observed
Fenno2
OB

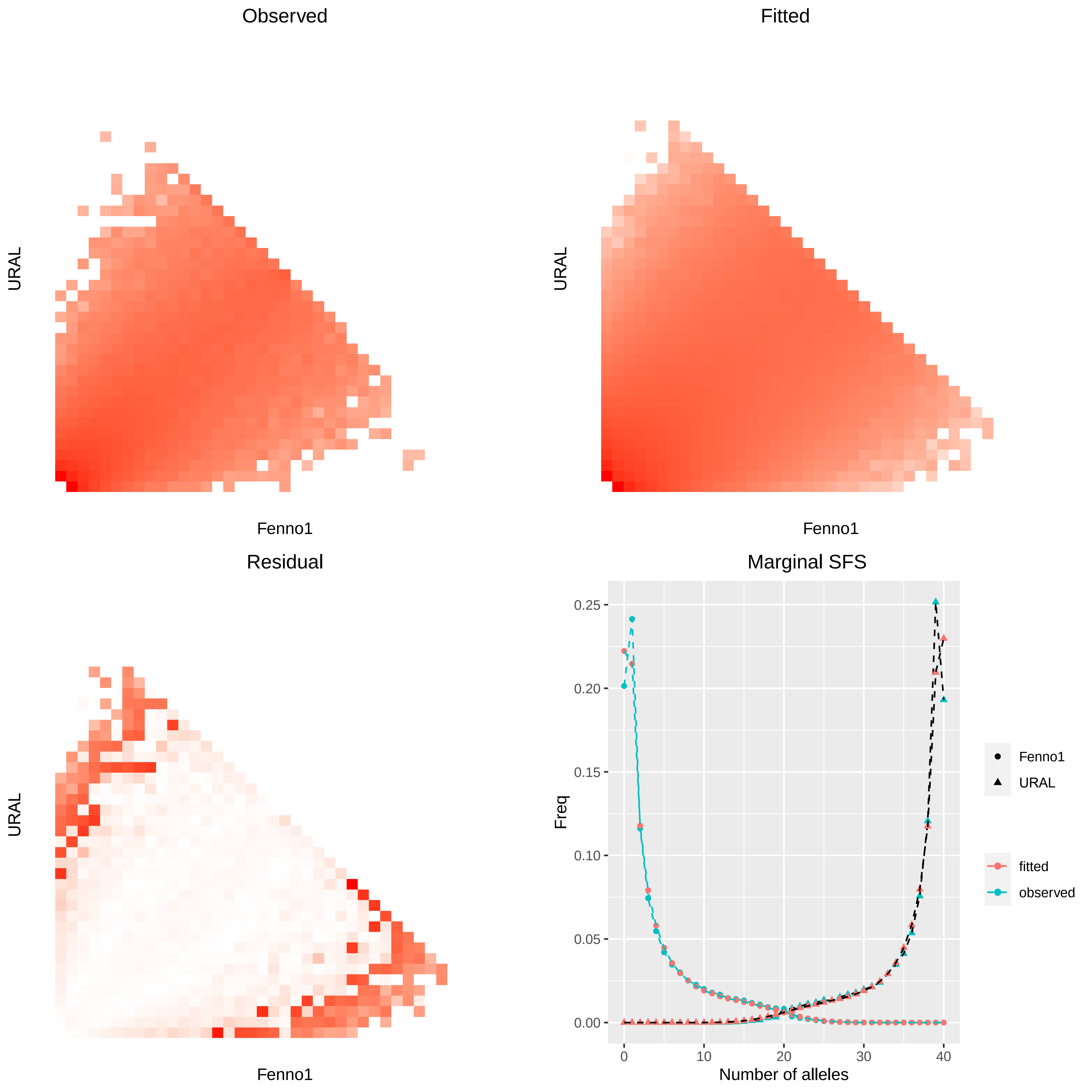
Observed
URAL
Fenno1
Fitted
URAL
Fenno1
Residual
URAL
Fenno1
Marginal SFS
0.25
0.20
0.15
0.10
0.05
0.00
Freq
0
10
20
30
40
Number of alleles
Fenno1
URAL
fitted
observed

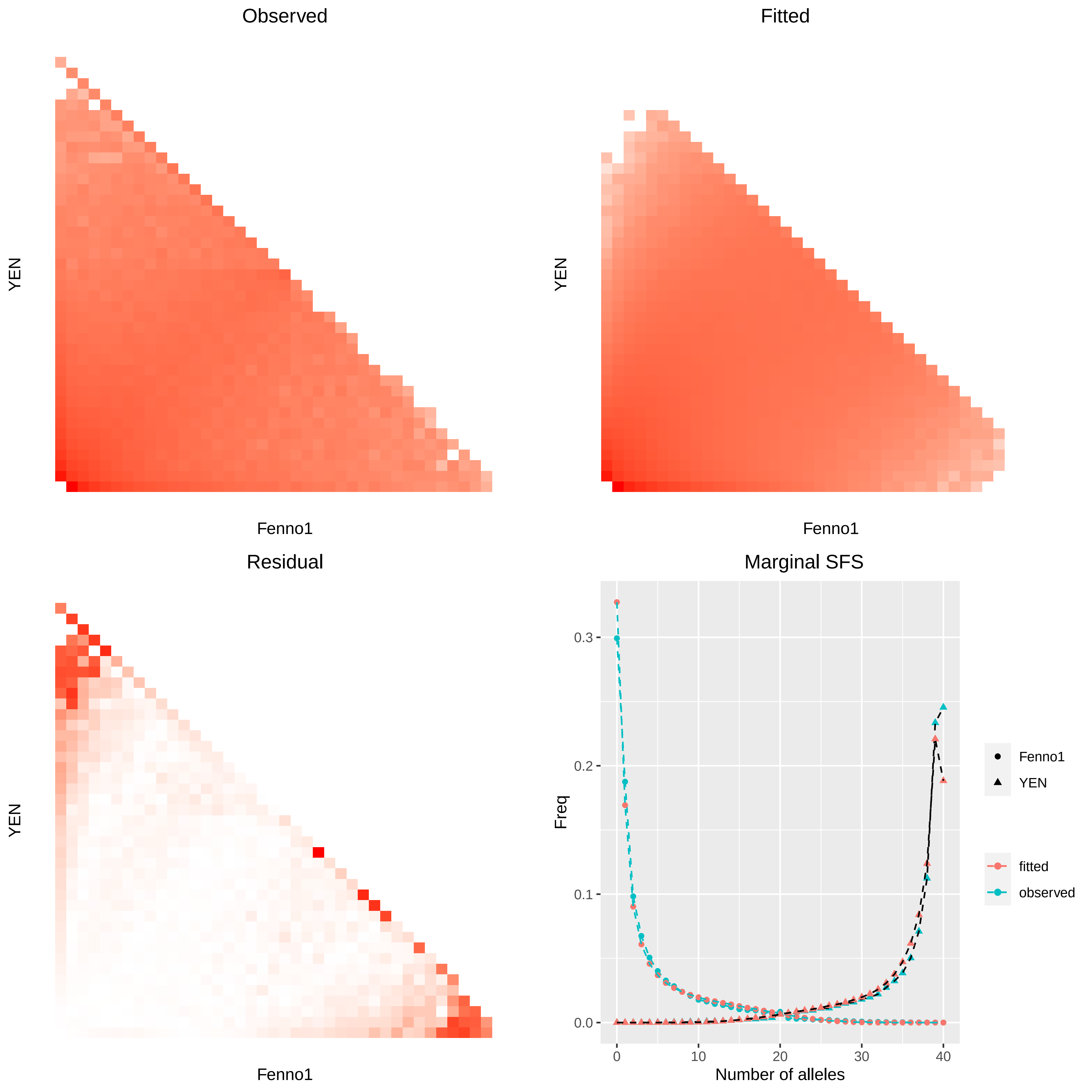

Observed
YEN
Fenno1
Fitted
YEN
Fenno1
Residual
YEN
Fenno1
Marginal SFS
Freq
0.0
0.1
0.2
0.3
0
10
20
30
40
Number of alleles
Fenno1
YEN
fitted
observed

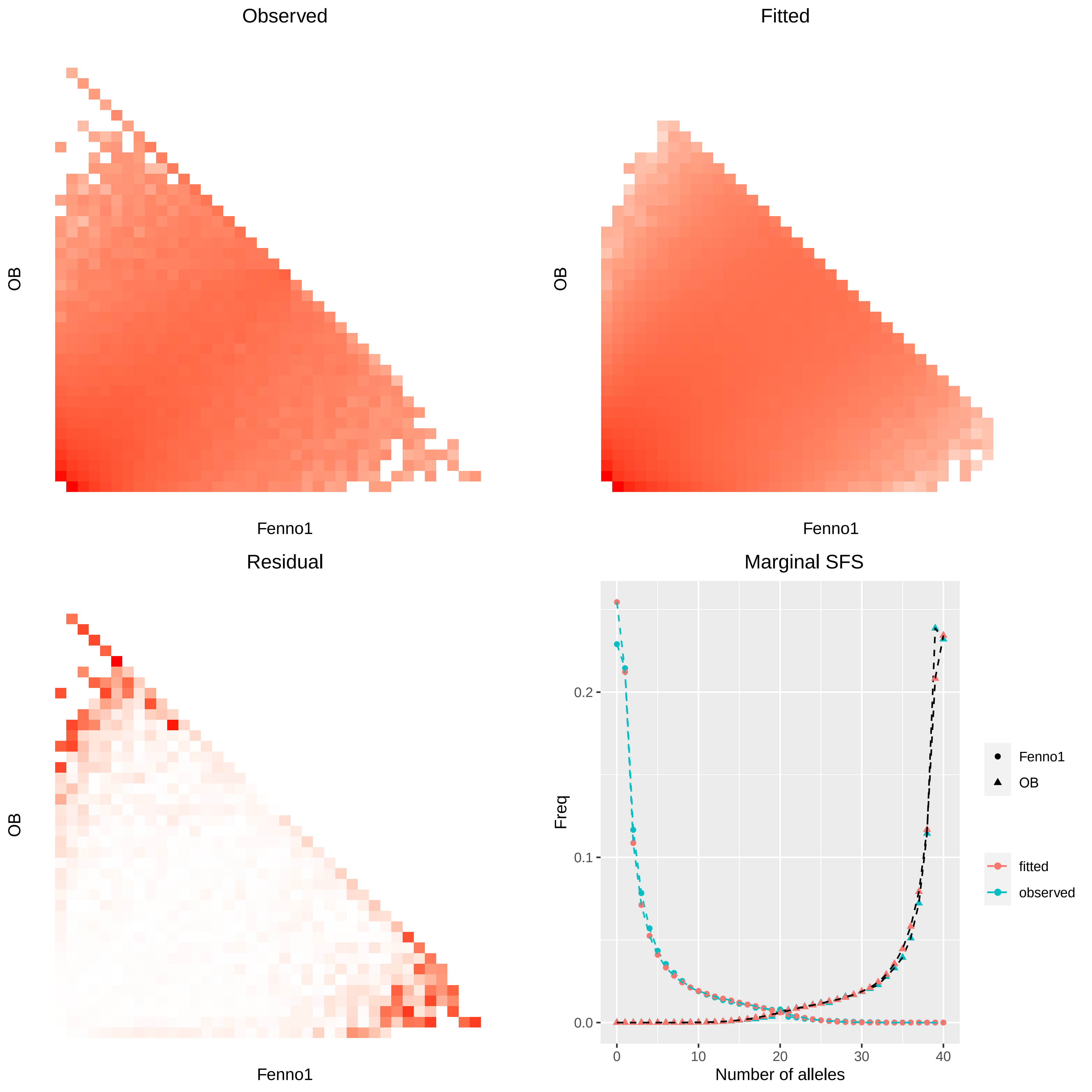
Observed
OB
Fenno1
Fitted
OB
Fenno1
Residual
OB
Fenno1
Marginal SFS
Freq
0.0
0.1
0.2
0
10
20
30
40
Number of alleles
Fenno1
OB
fitted
observed

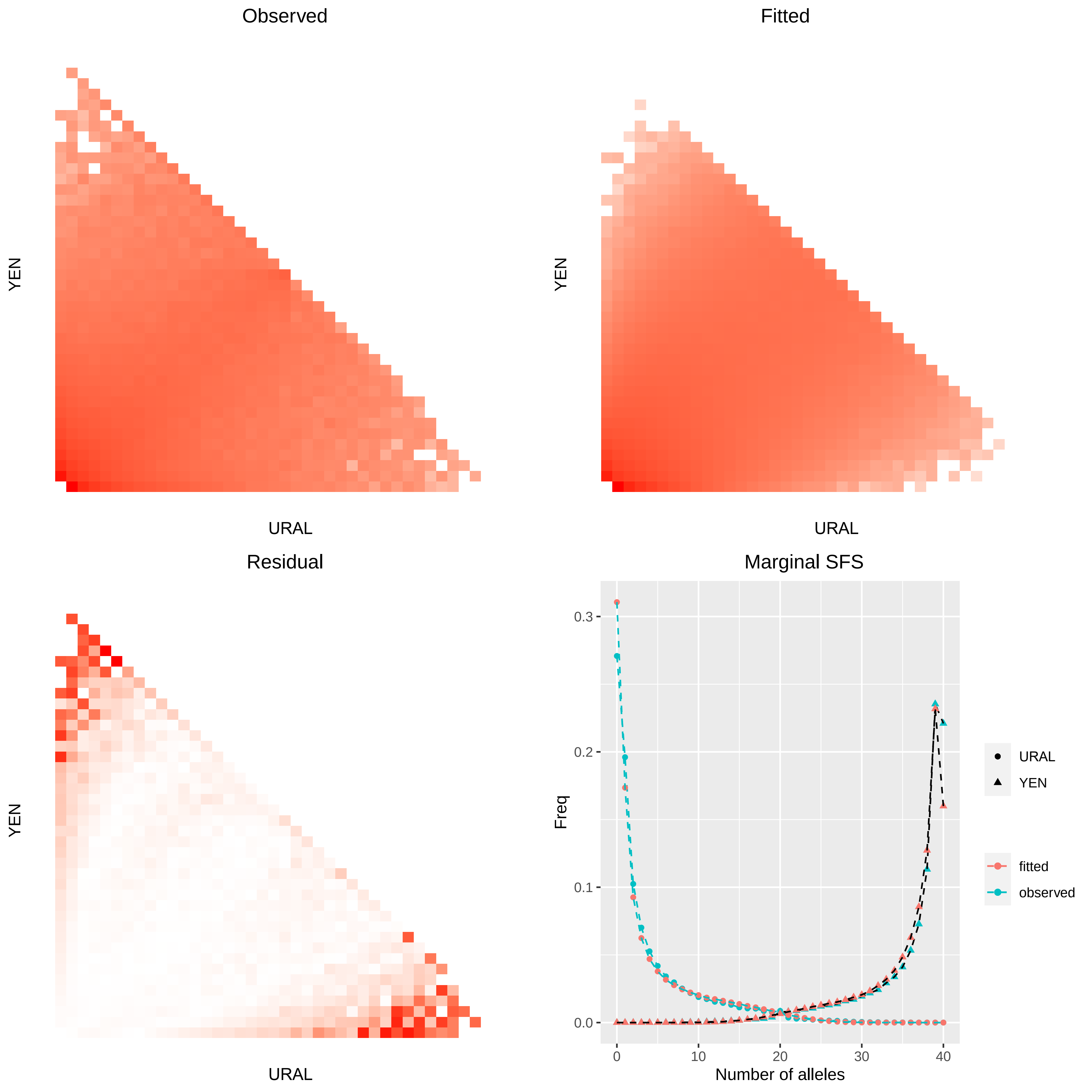
Observed
YEN
URAL
Fitted
YEN
URAL
Residual
YEN
URAL
Marginal SFS
Freq
0.0
0.1
0.2
0.3
0
10
20
30
40
Number of alleles
URAL
YEN
fitted
observed

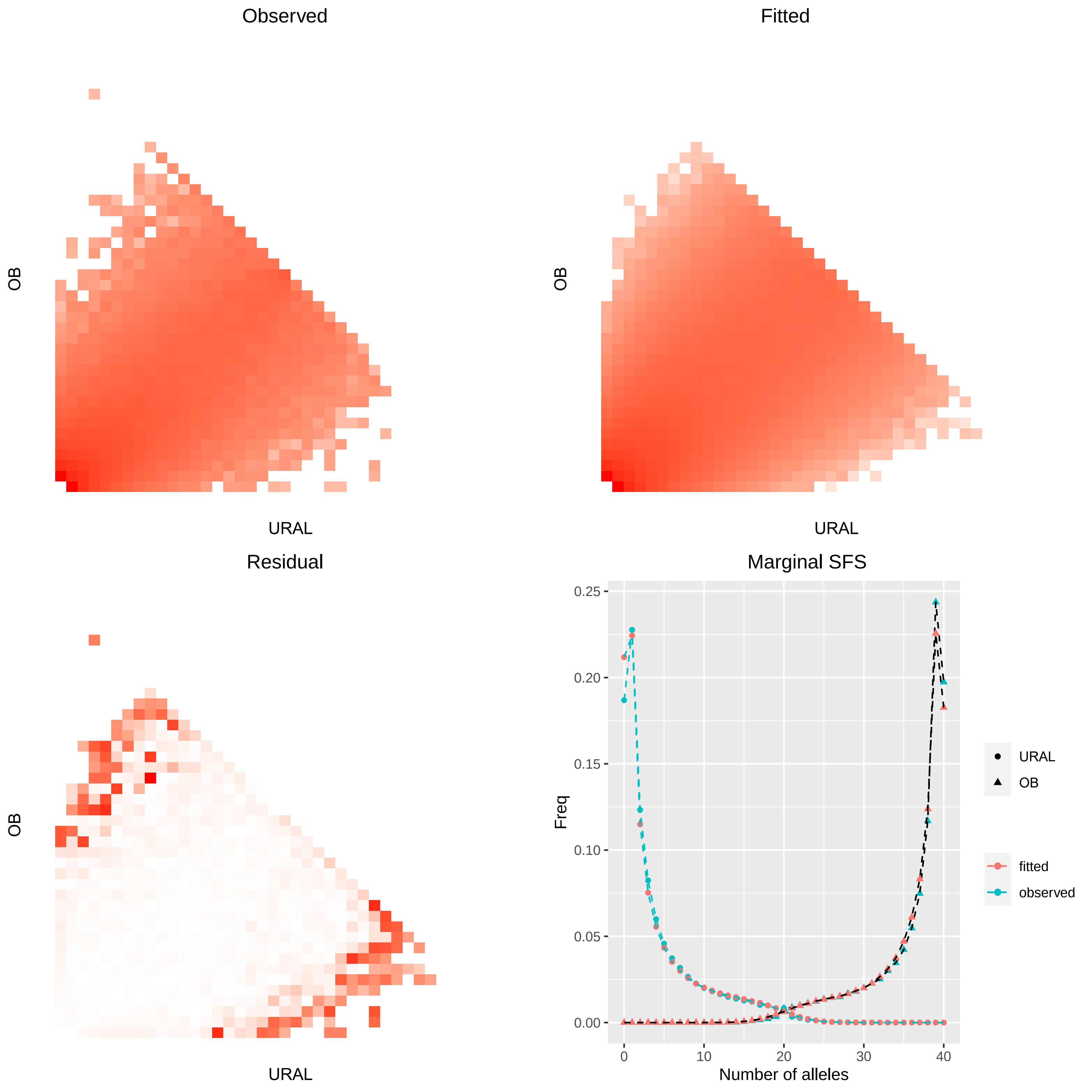
Observed
Fitted
OB
URAL
Residual
Marginal SFS
Freq
0.25
0.20
0.15
0.10
0.05
0.00
0
10
20
30
40
Number of alleles
URAL
OB
fitted
observed

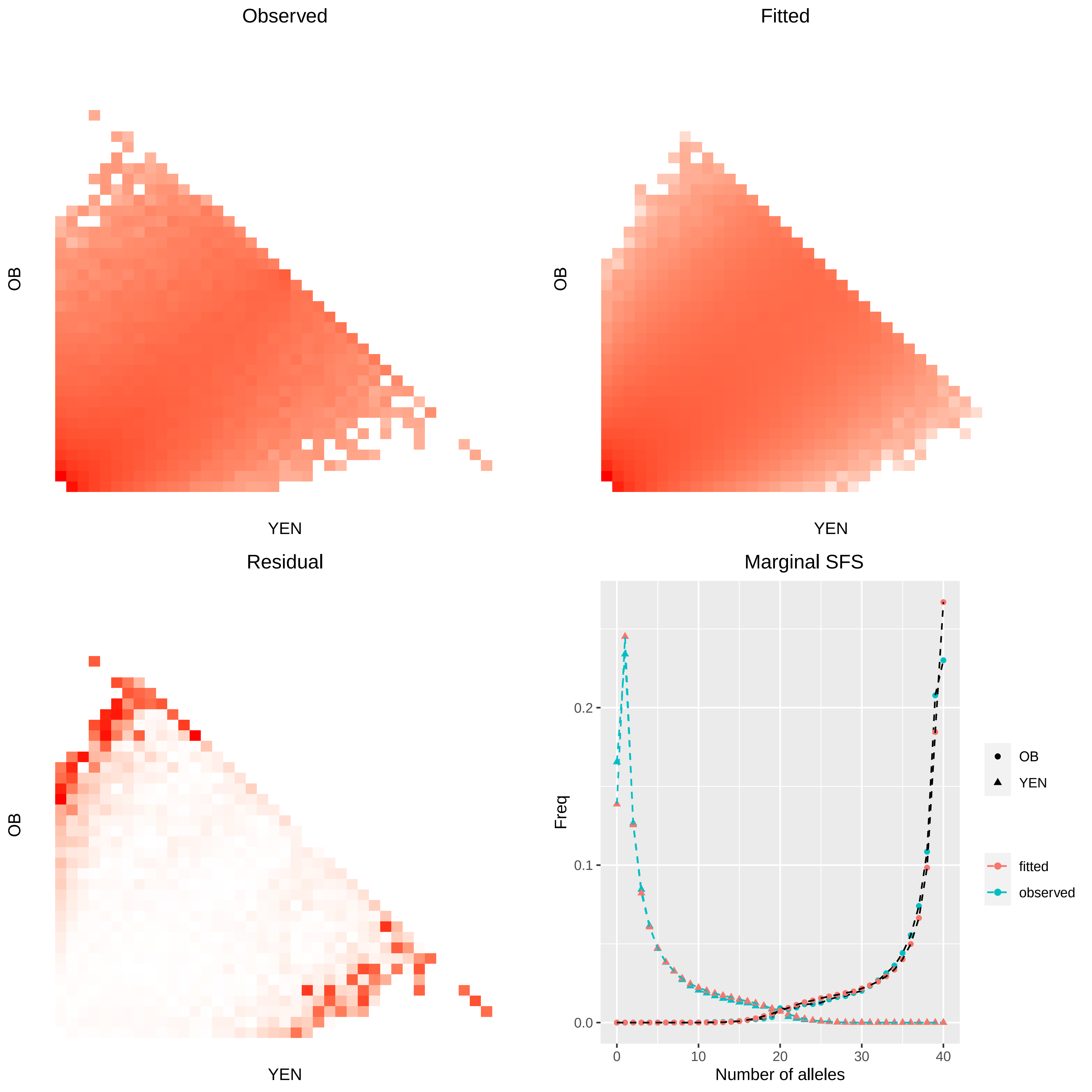

Observed
Fitted
Residual
Marginal SFS
OB
YEN
Freq
Number of alleles
0.0
0.1
0.2
0
10
20
30
40
OB
YEN
fitted
observed
